# Scalable empirical mixture models that account for across-site compositional heterogeneity

**DOI:** 10.1101/794263

**Authors:** Dominik Schrempf, Nicolas Lartillot, Gergely Szöllősi

## Abstract

Biochemical demands constrain the range of amino acids acceptable at specific sites resulting in across-site compositional heterogeneity of the amino acid replacement process. Phylogenetic models that disregard this heterogeneity are prone to systematic errors, which can lead to severe long branch attraction artifacts. State-of-the-art models accounting for across-site compositional heterogeneity include the CAT model, which is computationally expensive, and empirical distribution mixture models estimated via maximum likelihood (C10 to C60 models). Here, we present a new, scalable method EDCluster for finding empirical distribution mixture models involving a simple cluster analysis. The cluster analysis utilizes specific coordinate transformations which allow the detection of specialized amino acid distributions either from curated databases, or from the alignment at hand. We apply EDCluster to the HOGENOM and HSSP databases in order to provide universal distribution mixture (UDM) models comprising up to 4096 components. Detailed analyses of the UDM models demonstrate the removal of various long branch attraction artifacts and improved performance compared to the C10 to C60 models. Ready-to-use implementations of the UDM models are provided for three established software packages (IQ-TREE, Phylobayes, and RevBayes).

## 1 Introduction

Statistical uncertainty of phylogenetic analyses can be arbitrarily reduced by including more sequence data, which is today readily available given modern sequencing technologies. As a result, phylogenomic analyses based on complete genomes routinely provide very strong statistical support even for deep phylogenetic relationships. Statistical support, however, measures uncertainty in estimates assuming a specific evolutionary model and not accuracy of inferred phylogenetic inferences. Analyzing more sequence data alone cannot mitigate systematic biases that result from model misspecification or model inadequacy. In fact, more data can lead to arbitrary strong support for erroneous relationships under the wrong model (Philippe et al., 2011).

Long branch attraction (LBA) is a systematic bias in phylogenetic inference where branches are estimated to be shorter than they actually are (Felsenstein, 1978; Philippe et al., 1998). LBA may result in topological errors, and distantly related species may appear to be closer related. LBA artifacts are especially abundant, when inferring phylogenies using maximum parsimony, where multiple character changes are disregarded. The development of substitution models (Jukes et al., 1969) accounting for the possibility of multiple character changes has decreased the severity of LBA artifacts, especially when accounting for rate heterogeneity, for example with a discrete Gamma probability distribution (Yang, 1994b). In the following, the term *distribution* is used to refer to probability distributions.

Classical substitution models assume that sites in the sequence alignment of interest evolve according to a transition rate matrix describing the rates of change between different pairs of characters. The transition rate matrix is parametrized by a set of exchangeabilities between characters and a stationary distribution of characters. Usually, a single transition rate matrix is used for the entire alignment, and exchangeabilities and the stationary distribution are shared across all sites. Most often, the stationary distribution is set to the distribution of observed characters in the analyzed alignment. For alignments of amino acid sequences of real-life proteins, however, a shared stationary distribution across sites is clearly not adequate, because biochemical constraints limit the range of amino acids acceptable at specific sites reducing amino acid diversity in a site-specific manner (Pál et al., 2006; Goldstein, 2008; Franzosa et al., 2008). For example, at a specific site, an amino acid with a specific hydrophobicity, size, or mass may be required.

Phylogenetic inferences with models that disregard heterogeneity in the stationary distribution across sites (*across-site compositional heterogeneity*), have led to strongly supported LBA artifacts (Williams et al., 2013; Simion et al., 2017; Feuda et al., 2017). One reason for the underestimation of the lengths of long branches is that when only a reduced set of amino acids is used, the substitution process becomes saturated earlier than when the full set of amino acids is employed. This happens, because the probability of observing the same amino acid increases if the stationary distribution is constrained to a strict subset of all amino acids. Models that ignore a variation in the stationary distribution across sites, and instead use an averaged stationary distribution, will systematically underestimate the probability of observing the same amino acid, and consequently underestimate the branch length between two evolutionary distant observations. In phylogenetic terms this corresponds to a systematic underestimation of the probability of homoplasy (independent substitution events leading to the same amino acid) which can result in long branches being attracted because identical amino acid characters are erroneously interpreted as synapomorphies (i.e., resulting from a single substitution on an ancestral branch).

Across-site compositional heterogeneity has been modeled using partition models (e.g., Lanfear et al., 2016), and mixture models, the focal point of this contribution. In addition, mixture models of full transition rate matrices have been examined (Le et al., 2008b; Le et al., 2010; Le et al., 2012). Mixtures of full transition rate matrices allow different sites not only to exhibit specific amino acid compositions, but also to evolve with different exchangeabilities according to solvent exposure, protein structure or protein function.

In contrast to the above, the CAT model (Lartillot et al., 2004) uses one set of exchangeabilities for all mixture model components and a Dirichlet process prior over the stationary distributions and their weights in a Bayesian Markov chain Monte Carlo framework. The CAT model is widely used, and greatly improves model fit. However, computational requirements are high, such that convergence times are long, and convergence may be beyond reach, especially for larger data sets (Whelan et al., 2016).

Inspired by the CAT model, and consistent with the derivation of widely used empirical transition rate matrices from curated databases, empirical stationary distribution mixture (EDM) models, which use a fixed set of stationary distributions, have been developed. The rationale is that site-specific amino acid constraints may be caused by universal biochemical constraints (e.g., Jimenez et al., 2018). In particular, composition heterogeneity and site-specific amino acid constraints have already been used to estimate protein structure (Goldman et al., 1996) and the association of protein structure with evolution (Goldman et al., 1998).

Previously, Quang et al. (2008) used an expectation maximization algorithm to find EDM models with 10, 20, up to 60 components, which we collectively call CXX models, from alignments of the HSSP database (Schneider et al., 1997). Each mixture model component is defined by the used stationary distribution and weight. Accordingly, we use the term *component* to refer to a stationary distribution with corresponding weight. For computational reasons, the Poisson model (Felsenstein, 1981), which exhibits uniform exchangeabilities, was used when searching for the components. The phylogeny for each alignment was estimated beforehand using the WAG model. In contrast, Wang et al. (2008) use principal component analysis to detect four stationary distributions of amino acids from alignments of the Pfam data base (Sonnhammer et al., 1997). Inferences with EDM models such as the CXX models are much less computationally expansive than with the CAT model because they can be used in a maximum likelihood framework, where they exhibit good statistical fit.

Recently, a composite likelihood approach was developed that estimates stationary distributions of amino acids directly from the data at hand (Susko et al., 2018). Special strategies to estimate the stationary distributions need to be used, because if species are closely related, the observed amino acids are expected to be more similar. These strategies include (1) restricting the analysis to sites with high rate, (2) penalizing low frequencies of amino acids, (3) down-weighing contributions from species-rich clades, and (4) phylogeny-based estimation.

Here, we describe EDCluster, a new method for obtaining stationary distributions that can be used to construct EDM models. EDCluster can be used on any set of alignments ranging from large databases of homologous genes to more specific data sets. We employ the CAT model implemented in Phylobayes (Lartillot et al., 2013) to estimate site-specific posterior distributions of the stationary distributions of amino acids. In this way, specialized treatment of the expected variation in the divergence between the sequences is not required. The site distributions are analyzed as is, or transformed using linear transformations developed specifically for compositional data. The transformations aid the clustering method in finding stationary distributions of amino acids with different specialized features. The use of a clustering algorithm seemed natural because clustering is a simple machine learning approach for feature discovery. Although EDCluster does not directly use biochemical information, the inferred components are found to correspond to specific biochemical traits of amino acids, such as hydrophobicity, size, or mass.

Using EDCluster, we provide sets of 4, 8, 16, …, 4096 components estimated from subsets of the HOGENOM database (Dufayard et al., 2005), the HSSP database, and the union of both. We present extensive analyses of EDM models based on these sets of components which we collectively call universal distribution mixture (UDM) models. For the same number of components, we demonstrate that the UDM models outperform the CXX models not only in terms of likelihood but also in parametric bootstrap analyses, were they exhibit improved amino acid compositions and branch lengths. Moreover, EDCluster allows construction of EDM models with a large number of components. In particular, the UDM models with more components show even further increases in accuracy. However, the number of components is still limited by the associated linear increase of computational requirements during inference. In conclusion, the UDM models minimize systematic errors caused by constraints in amino acid usage in a fraction of the run time of CAT. We provide ready-to-use implementations for several established phylogenetic software packages such as IQ-TREE (Nguyen et al., 2015), Phylobayes, or RevBayes (Höhna et al., 2016). Further, we provide user-friendly scripts implementing EDCluster to construct EDM models specific to the data set at hand. Finally, we employ a simulation study to reproduce a well known LBA artifact of classical substitution models, and show that application of EDM models successfully recovers the correct topology.

## 2 New Approaches

EDM models assume that evolution occurs along a phylogeny according to a mixture of *N* amino acid substitution models. The transition rate matrices of the different components share a single set of exchangeabilities. In this contribution, Poisson exchangeabilities were used exclusively although the method could in principle be generalized directly to any other set of exchangeabilities. In contrast, the stationary distributions (or equilibrium frequencies over the 20 amino acids) differ between each component of the mixture model. Here, the stationary distributions are inferred from alignments obtained from curated databases. Each alignment was analyzed with Phylobayes under the CAT model with Poisson exchangeabilities. For each alignment and each site, the expectation of the posterior distribution of the stationary distribution of amino acids (*site distribution*), was calculated. Each site distribution is a point in 20-dimensional space with elements summing up to 1.0. For each database, the site distributions of all sites were used as is or transformed before further analysis. Application of linear transformations is a standard procedure in analyses of compositional data. The two employed transformations were: (1) the centered log ratio transformation (CLR; Aitchison, 1982), and (2) the log centered log ratio transformation (LCLR; Godichon-Baggioni et al., 2018). In our case, the transformations ensure that site distributions exhibiting specific different features are moved further apart from each other, so that they fall into different groups in the subsequent analysis. *K*-means clustering was used to group the prepared site distributions into *N* ∈ {4, 8, 16, …, 4096} clusters. The stationary distributions and weights of the different components to be used in the UDM models were set to the determined cluster centers and their relative weights, respectively.

We combine the components obtained from a subset of the HOGENOM database with Poisson exchangeabilities and refer to the mixture model resulting from a specific set of components as UDM-XXX-Trans, where XXX is the number of components, and Trans is the used transformation (None, CLR, or LCLR). The usage of Poisson exchangeabilities is implicitly assumed and not mentioned specifically. For example, the UDM model with four components obtained from the LCLR transformed site distributions is referred to as UDM-004-LCLR model. Although the presented analyses exclusively focus on components estimated from the HOGENOM database, components estimated from the HSSP database, and the union of both databases are provided for further reference.

## 3 Results

### Analysis of UDM model components

EDM models differ by their used set of stationary distributions and weights (components). The effective number of amino acids (*K*_eff_, see Material and Methods) measures the diversity of discrete distributions. For stationary distributions of amino acids, *K*_eff_ values range from 1 for highly constrained stationary distributions with only one used amino acid to 20 for the uniform stationary distribution of amino acids which is used by the Poisson model. Most often, the empirical distribution of amino acids observed in the analyzed alignment is used for inference. Usually, these empirical stationary distributions exhibit a high effective number of amino acids of 15 < *K*_eff_ < 20. In particular, the default stationary distribution of the LG model (Le et al., 2008a) has *K*_eff_ = 18.04.

The performance of an EDM model is strongly characterized by the composition of effective number of amino acids of the used stationary distributions together with their weights. In general, UDM models with more components employ more specialized, constrained stationary distributions with lower *K*_eff_ values, and also put more weight on these constrained distributions (Figure 1). Accordingly, the mean *K*_eff_ value decreases with the number of components. In particular, a general, “catch-all” stationary distribution exhibiting *K*_eff_ ≈ 17 is retained. The weight of the most general stationary distribution decreases with the number of components. Additional components exhibit stationary distributions with *K*_eff_ values usually well below 10. UDM models with more than 128 components tend to include more than one general stationary distribution with *K*_eff_ > 10. The results for the stationary distributions obtained from untransformed, and CLR transformed site distributions are almost identical (Figure S1 and Figure S3).

**Figure 1:**
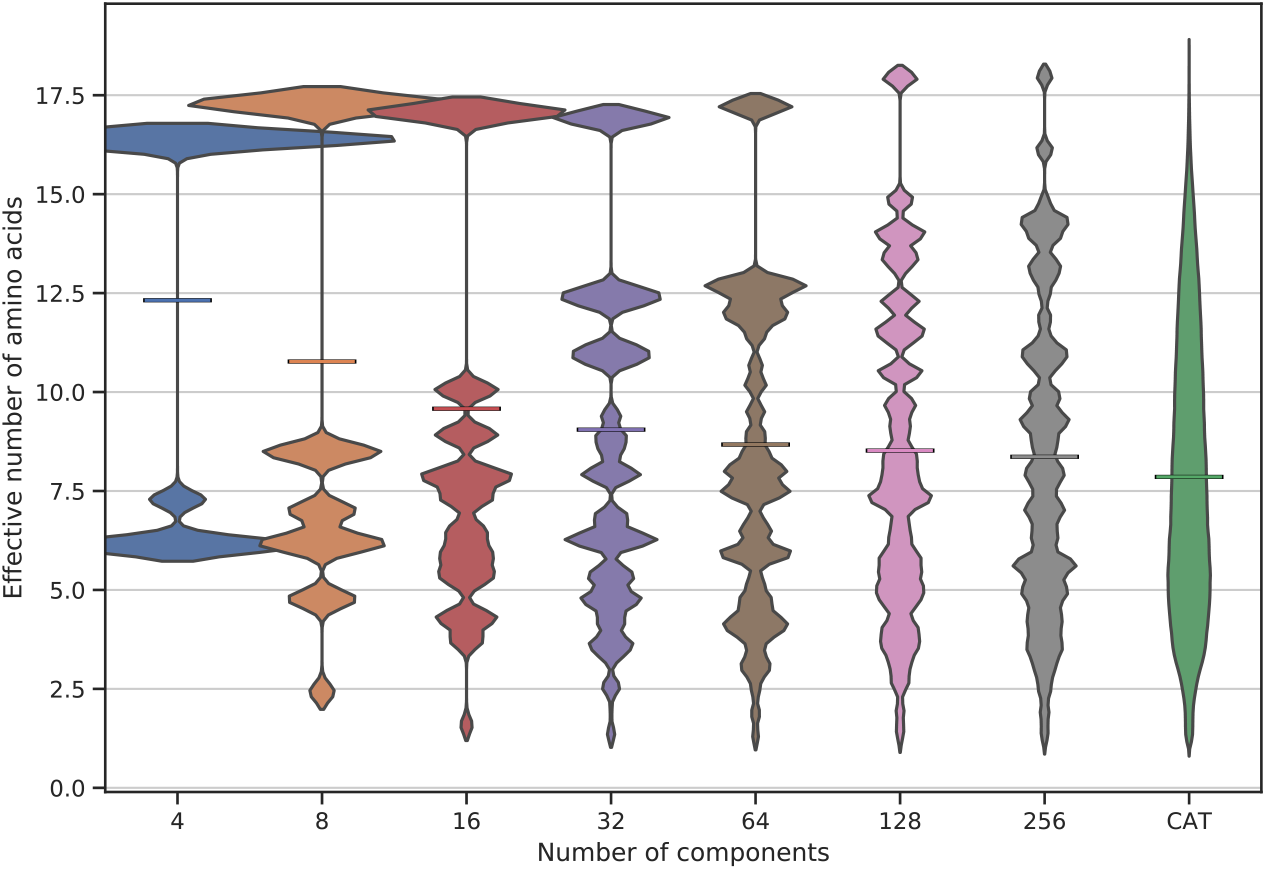
Distributions of effective number of amino acids of the stationary distributions used by universal distribution mixture models with different numbers of components. Violin plots of the effective number of amino acids of the stationary distributions obtained from the log center log ratio transformed site distributions (Godichon-Baggioni et al., 2018) of the HOGENOM database (Dufayard et al., 2005) are shown. On the far right, the distribution of the effective number of amino acids of the site distributions obtained with the CAT model (Lartillot et al., 2004) using Poisson exchangeabilities (Felsenstein, 1981) is displayed. The width of the violin plots was normalized such that all areas are equal. Horizontal bars display the means of the distributions.

Further, the distribution of *K*_eff_ values of the site distributions inferred from the HOGENOM database using the CAT model with Poisson exchangeabilities and the corresponding mean value are shown. As expected, the more components are used, the closer the distribution of *K*_eff_ values of the stationary distributions of the UDM components to the distribution estimated directly from the HOGENOM database (see also Figures S2, S4, and S5). The mean *K*_eff_ values exhibit the same tendency. Exact *K*_eff_ values for UDM models up to 16 components including all three transformations are available in Section S3. Strikingly, in most cases components with higher weight also have higher *K*_eff_ values.

Further, the *K*_eff_ values of the site distributions associated with the components with the most general stationary distributions are usually much lower than the *K*_eff_ value of the stationary distribution of the respective component. In particular, the first component sorted by weight of the UDM-016-LCLR model exhibits a stationary distribution with *K*_eff_ = 17.1, but the median of the *K*_eff_ values of the associated site distributions is 12.2 (Figure 2, left; see Figure S6 for more details). Even for the UDM-256-LCLR model the first components exhibit a striking discrepancy between *K*_eff_ values (Figures S7 and S8). The substantial difference between the *K*_eff_ values of the site distributions and the stationary distribution of the corresponding component is only apparent for the first few components when sorting them according to weight. For example, the *K*_eff_ value of the stationary distribution of the fourth component (purple in Figure 2) is already very close to the median of the *K*_eff_ values of the site distributions. The components with lower weight show even higher agreement between the median *K*_eff_ value and the *K*_eff_ value of the cluster center. A more detailed analysis shows that the mean of the differences between the *K*_eff_ values of the site distributions and their associated cluster centers, which represents the loss in amino-acid specificity resulting from using a finite mixture of a given number of components, decreases monotonically with the number of components (Figure S9).

**Figure 2:**
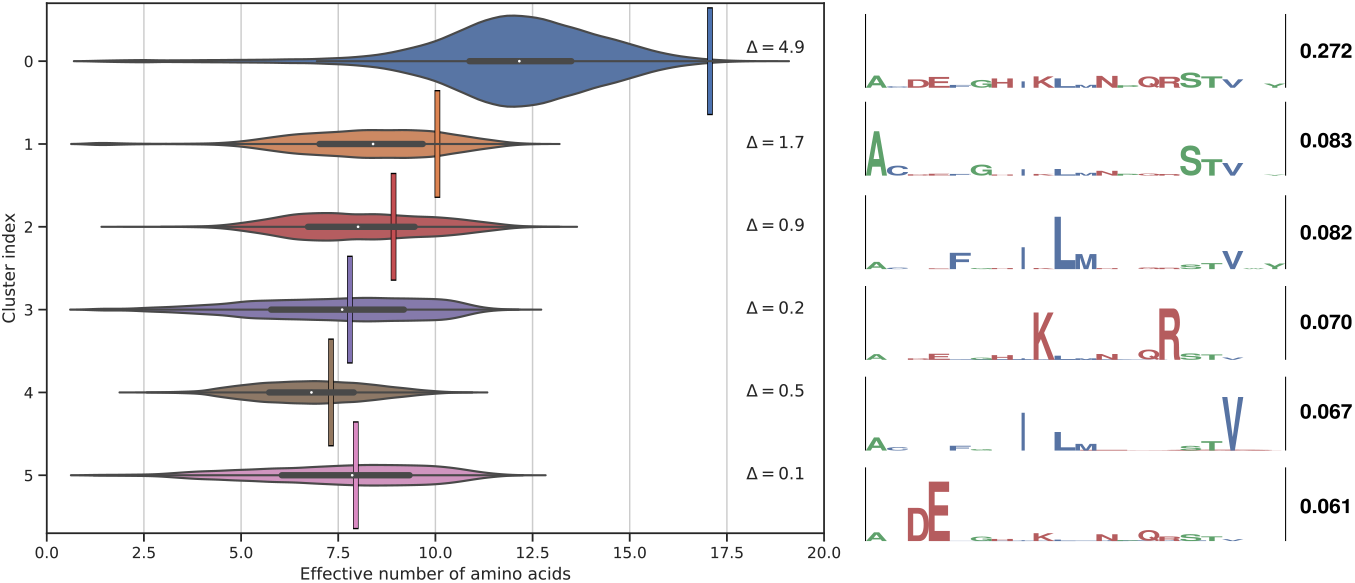
Analysis of the components of a universal distribution mixture (UDM) model. (Left) Violin plot of the effective number of amino acids of the site distributions associated with the first six components sorted by weight of the UDM model with a total number of 16 components obtained from clustering the log center log ratio (LCLR; Godichon-Baggioni et al., 2018) transformed site distributions (UDM-016-LCLR model). The effective numbers of amino acids of the stationary distribution of the components themselves are shown by vertical lines. The respective differences to the medians of the associated site distributions are also given (Δ values). (Right) Customized WebLogos (Crooks, 2004) visualize general features of the amino acid distributions of the components. The heights of the amino acid letter codes correspond to their probabilities; the total height of each logo is 0.5. The amino acids are colored according to their hydrophobicity. Hydrophilic amino acids D, E, H, K, N, Q, and R, with hydrophobicity indices below −1.9 are colored in red. Hydrophobic amino acids C, F, I, L, M, and V, with hydrophobicity indices above 1.9 are colored in blue. Finally, amino acids A, G, P, S, T, W, and Y, with hydrophobicity indices between −1.9 and 1.9 are colored in green. The weights of the components are shown to the right of the logos.

Observe that the stationary distribution of the component with highest weight is very general (Figure 2), the second component is enriched for neutral amino acids with hydrophobicity indices close to zero, and the third and fourth component select for hydrophobic and hydrophilic amino acids, respectively. Also note that the weight of the first component differs significantly from the weight of the other five shown components. Altogether, the stationary distributions of the different first components of the UDM-016-LCLR model exhibit limited overlap and no apparent redundancy.

### Performance of UDM models

The performance of the UDM models was assayed on three empirical data sets that exhibit well characterized LBA artifacts when applying classical substitution models such as the LG model. The first data set encompasses eukaryotes including the fast evolving microsporidia and a distant archaeal outgroup. Microsporidia are a group of spore-forming unicellular parasites, which notably lack mitochondria (Cavalier-Smith, 1987). The lack of mitochondria and phylogenetic placement as the first emerging eukaryotic group (Vossbrinck et al., 1987; Kamaishi et al., 1996) marked them as a candidate for an ancient eukaryotic lineage predating the acquisition of mitochondria. However, more sophisticated phylogenetic analyses have recovered microsporidia being relatives of fungi, rather than being basal eukaryotes (Hirt et al., 1999; Keeling et al., 2000; Van de Peer et al., 2000; Keeling et al., 2002) and subsequently remnants of mitochondria were found experimentally (Williams et al., 2002). Here, as an illustration, we consider the data set of Brinkmann et al. (2005, referred to as microsporidia data set), which spans 40 sequences, and comprises 133 genes corresponding to 24294 amino acid sites. We show analyses of the concatenated alignment, as well as of the separate genes. For the concatenated alignment of the microsporidia data set, site homogeneous substitution models such as the LG model favor the former topology in terms of likelihood. Models accounting for across-site compositional heterogeneity such as the CAT model exhibit higher posterior probabilities for the latter topology, which is now widely accepted. Note that we use the term topology when referring to the order of branching events, and the term phylogeny, when referring to the topology together with branch lengths.

Further two data sets involving the positioning of nematodes and platyhelminths were analyzed (referred to as the nematode data set and the platyhelminth data set, respectively; Philippe et al., 2005). These data sets contain a total of 37, and 32 taxa with 35371 amino acid sites, respectively. The LBA artifacts, observed when using classical substitution models such as the LG model, are: nematodes and platyhelminths branching with a clade containing both, deuterostomes and arthropodes. Current phylogenetic consensus has nematodes and platyhelminths branching with arthropodes — a result strongly supported by the CAT model. In the following, we refer to the three topologies most likely exhibiting LBA artifacts as T1, and to the topologies in agreement with current phylogenetic consensus as T2 (Figure 3).

**Figure 3:**
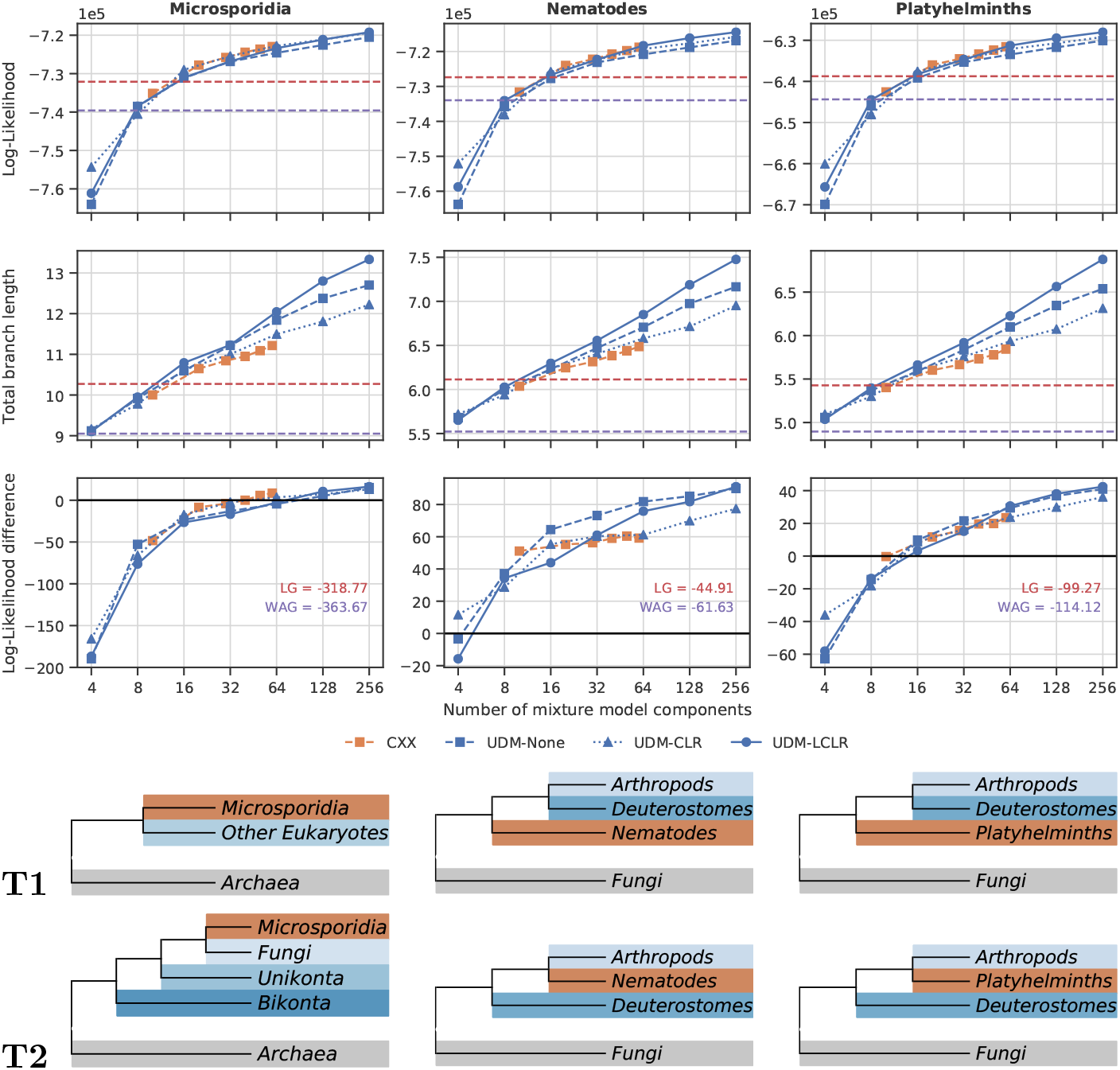
Performance of universal distribution mixture models (UDM; blue), and CXX models (orange; Quang et al., 2008) for an increasing number of components for the three empirical data sets. Results for UDM models are shown for the untransformed (None), center log ratio transformed (CLR; Aitchison, 1982), and log center log ratio transformed (LCLR; Godichon-Baggioni et al., 2018) site distributions. Results for WAG (purple; Whelan et al., 2001), and LG (red; Le et al., 2008a) are indicated by dashed horizontal lines. The rows from top to bottom show: (1) the maximum log-likelihoods, (2) the sum of all branch lengths (total branch length) of the maximum likelihood phylogenies measured in average number of substitutions, (3) the log likelihood differences between the two topologies presented below with a positive value indicating support for topology T2, (4) historical topology T1 affected by long branch attraction artifacts, and (5) currently accepted topology T2. In the topologies, outgroup, clade of interest, and ingroups are colored gray, red, and in shades of blue, respectively.

Maximum likelihood analyses were performed with IQ-TREE using UDM and CXX models with Poisson exchangeabilities, as well as the WAG and the LG model (Figure 3). Indeed, traditional substitution models favor the topologies T1 exhibiting the discussed LBA artifacts, whereas sufficiently component-rich UDM models reject T1 in favor of the currently accepted topologies T2 (Figures S10, S18, and S20). In general, the results agree very well across the three data sets. In terms of maximum log-likelihood, the WAG model performs slightly worse than the LG model. When using the same number of components, the maximum log-likelihood under the UDM and CXX models are similar. Eight and 16 components are needed to approximately achieve maximum log-likelihood values equivalent to the ones the WAG and LG models, respectively. Usage of more components further improves the maximum log-likelihood of the UDM and CXX models to the extent that they outperform classical substitution models, even-though they use Poisson exchangeabilities. The UDM models outperform the CXX models when using 64 components or more, because CXX models are not available with more than 60 components. Bayesian information criterion (BIC, Schwarz, 1978) scores are monotonically decreasing with the number of components, and component-rich models are clearly favored (Figures S11, S19, and S21).

For the UDM and CXX models, the total branch length of the maximum likelihood phylogenies increases with the number of used components. When increasing the number of components, the total branch lengths do not approach a limit but exhibit logarithmic increase. The total branch lengths of the maximum likelihood phylogenies of the WAG model are lower than the ones of the UDM model with four components. The total branch lengths of the phylogenies obtained by the LG model are surpassed when using eight to 16 components, approximately. The total branch lengths of the maximum likelihood phylogenies of the UDM models tend to be larger than the ones of the CXX models. The transformation affects total branch lengths more than the other presented results. Components obtained from the LCLR transformed site distributions exhibit highest total branch lengths.

Next, the power to discriminate between topologies T1 and T2 was examined. To this end, the maximum log-likelihoods of analyses constrained to the two different topologies T1 and T2 were compared. The topologies were fixed during the analyses, but the branch lengths and other model parameters were inferred. The difference of the maximum log-likelihood values acquired from topologies T2 and topologies T1 indicates whether the LBA artifacts are supported (negative values), or rejected (positive values). The WAG and LG models both strongly support the topology exhibiting the LBA artifacts in all three cases with large differences in maximum log-likelihood. In contrast, if the number of components is large enough, the UDM and CXX models reject the topology exhibiting the LBA artifacts in all three data sets. For the data set involving microsporidia, compared to the CXX models the UDM models require a higher number of components to reject the topology with LBA artifacts. For the data sets involving nematodes and platyhelminths, the situation is reversed in that the differences of the maximum log-likelihoods of the UDM models are more positive than the ones of the CXX models. Also, the difference in maximum log-likelihood does not increase substantially for the CXX models when applied to the data set involving nematodes.

The performance of the UDM models on shorter alignments compared to the LG model, and the CAT model with Poisson exchangeabilities was tested on the separate genes of the microsporidia data set. The inferred trees were compared to the historical topology T1 (Figures S12–S14) and the more recent topology T2 (Figures S15–S17). The lengths of the alignments range from 40 to 600 amino acid columns. Consequently, we observe high variance between the results from different genes. The means of the symmetric (Robinson et al., 1981) and branch score (Kuhner et al., 1994) distances to the T1 topology do not improve when using models accounting for across-site composition heterogeneity compared to the LG model. However, the means of the symmetric and branch score distances do decrease with the number of components for the T2 topology. The branch score distance of the CAT model is larger than for the UDM-128-LCLR. We stress that there is no strong consensus of the current phylogenetic literature about the branch lengths of the discussed tree. In general, the difference between the results of the UDM models and the CAT model decreases with the number of components. The results of the UDM-128-LCLR model and the CAT model are nearly identical. The incompatible split distance, which is a distance measure accounting for the uncertainty in the inferred topology, shows a consistent decrease of the distance with the number of components.

### Model adequacy in recovering across-site compositional heterogeneity

Finally, we assayed the potential of the UDM, and CXX models, as well as of the WAG and the LG models to reproduce the across-site compositional heterogeneity of empirical alignments. For this reason, we preformed parametric bootstrap in a manner similar to posterior predictive analyses in Bayesian statistics. We estimated model parameters using maximum likelihood for the microsporidia, nematode, and platyhelminth data sets and used these to simulate alignments comprising 25 000 sites, which is close to the length of the original data sets. Subsequently, summary statistics for the original alignments, and the simulated alignments were compared. It is desirable that the simulated alignments reproduce characteristics of the original alignments.

Here, we compared the distribution of effective number of amino acids observed at each site of the alignment. It is difficult to judge differences in the actual distributions of effective number of amino acids by eye (Figures S22, S23, and S24). Therefore, we present the Wasserstein distance (also known as earth mover’s distance) between the distributions of effective number of amino acids of the original and the simulated alignments.

For all three data sets, the WAG and LG models with a single amino acid transition rate matrix produce alignments with inflated diversity measured in effective number of amino acids (Figure 4 for the microsporidia data set; Figures S25, and S26 for the nematode and platyhelminth data sets, respectively). Numeric values of the average effective number of amino acids per site in the alignment, as well as the Wasserstein distances are given in Table S1. Component-rich UDM and CXX models typically exhibit lower Wasserstein distances than UDM or CXX models with fewer components. The UDM models consistently outperform the CXX models. The C50 model exhibits large deviations in all three data sets. UDM models with 256 components sometimes exhibit higher Wasserstein distances than their equivalents with 128 components. For example, compare the UDM-256-LCLR model with the UDM-128-LCLR model in the microsporidia data set (Figure 4). The reason for the increase of the Wasserstein distance is that the average number of used amino acids of the UDM-256-LCLR model is actually lower than the one of the original data.

**Figure 4:**
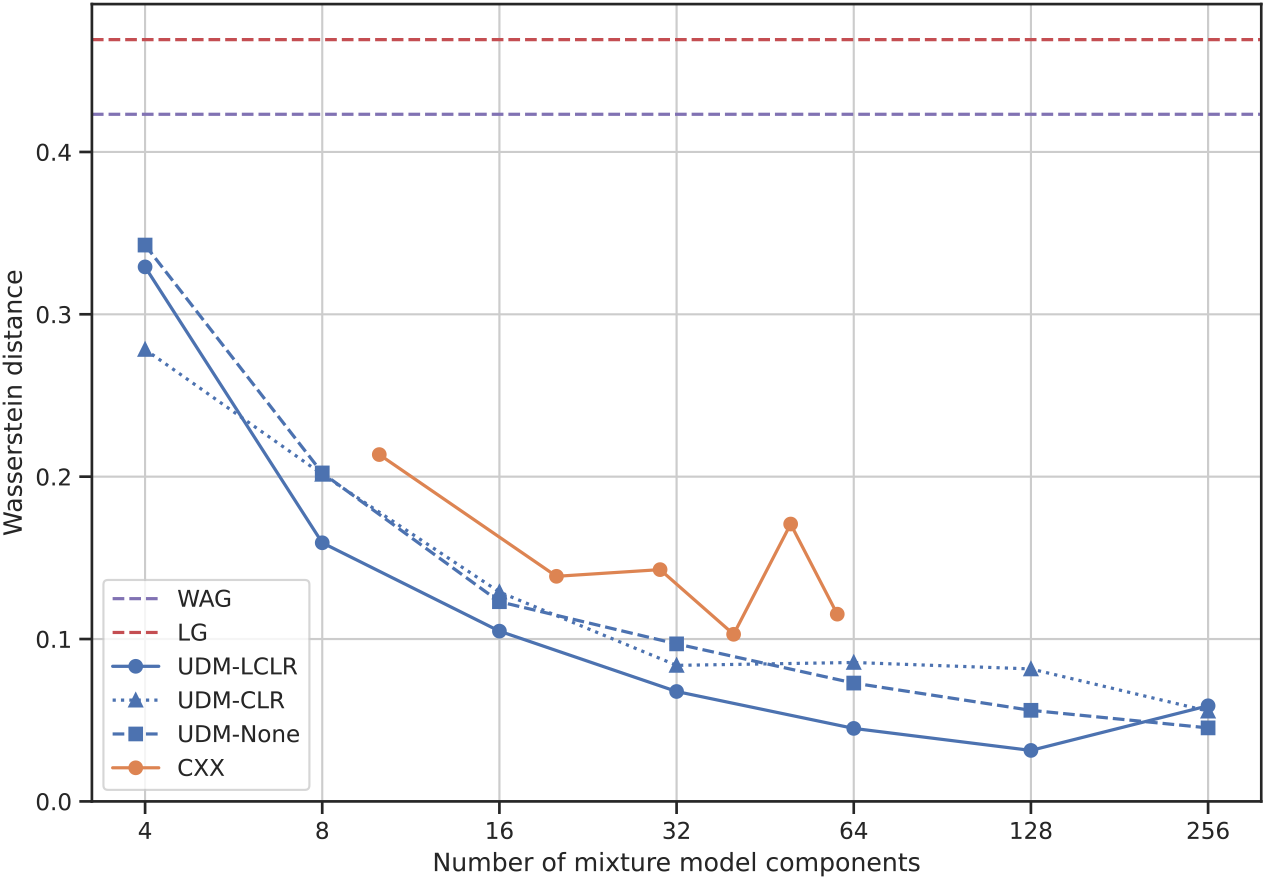
Across-site compositional heterogeneity of classical substitution models and empirical distribution mixture models. Similarity between the across-site compositional heterogeneity of the microsporidia data set (Brinkmann et al., 2005), and simulated alignments for the maximum likelihood parameter estimates of the WAG (Whelan et al., 2001), LG (Le et al., 2008a), CXX (Quang et al., 2008), and universal distribution mixture (UDM) models. Results of UDM models obtained from untransformed (None), center log ratio (CLR; Aitchison, 1982) transformed, and log center log ratio (LCLR; Godichon-Baggioni et al., 2018) transformed site distributions are shown. Similarity is measured by the Wasserstein distance between the distributions of effective number of amino acids per site between empirical data and the sequences simulated using parametric bootstrap.

### Phylogenetic artifact can be reproduced in simulation study

The results presented above rely on assumptions about the correct topology (Whelan et al., 2016). As an alternative, we can experiment with simulations, for which we know the true phylogeny. Interestingly, the LBA artifact observed in the microsporidia data set could be reproduced in a simple simulation study. We used the 175 330 site distributions obtained from the HOGENOM database to simulate an alignment along a phylogeny exhibiting the currently accepted topology T2 where microsporidia branch within fungi (left phylogeny in Figure 5). Then, maximum-likelihood phylogenies were inferred with the Poisson model, and the LG model, as well as the CXX models, and the UDM models which account for across-site compositional heterogeneity. The maximum-likelihood phylogenies of the Poisson and LG models exhibit the incorrect topology T1 where microsporidia are positioned at the eukaryotic root (Figures 5 and S27). In addition, the ciliates are also moved outside their clade. All branches are supported with bootstrap values of 100 %.

**Figure 5:**
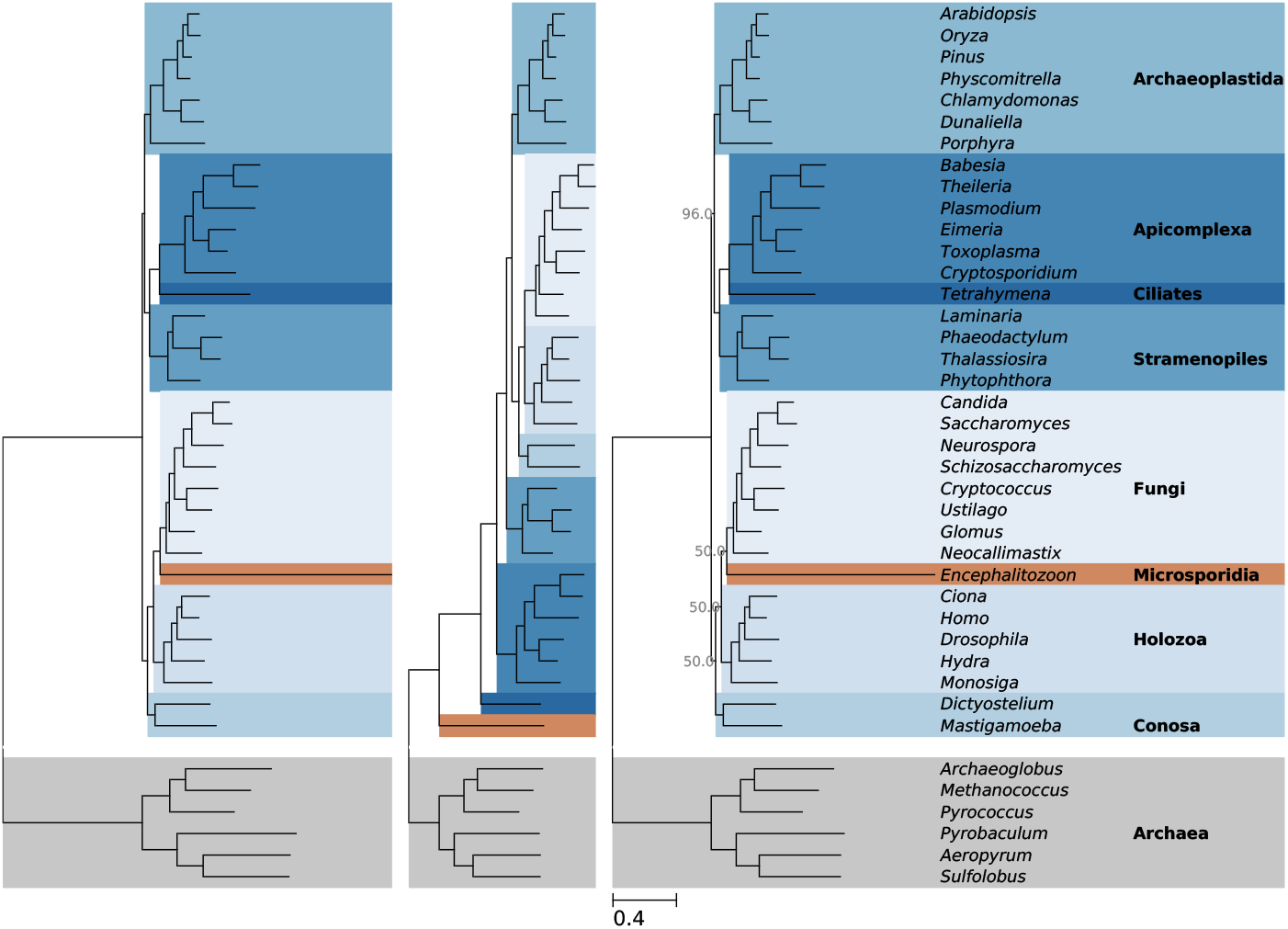
Reproduction of microsporidia long branch attraction artifact in simulation study. (Left) Phylogeny used for simulation using Poisson exchangeabilities (Felsenstein, 1981), and stationary distributions of amino acids obtained from analyses of the HOGENOM database (Dufayard et al., 2005). (Middle) Maximum likelihood phylogeny of the Poisson model. (Right) Maximum likelihood phylogeny of the universal distribution mixture model with 32 components obtained from log center log ratio transformed (Godichon-Baggioni et al., 2018) site distributions (UDM-032-LCLR model). For both inferred phylogenies, bootstrap values below 100 % are shown.

The maximum likelihood phylogenies inferred by UDM models with 4, 8 and 16 components still exhibit the LBA artifact involving microsporidia (Figure S28). In contrast, the UDM-032-LCLR model correctly supports microsporidia branching from within fungi. The correct phylogeny has much higher statistical support with an improvement in BIC score of 2591445 units compared to the results of the Poisson model. The position of microsporidia, holozoa, and conosa still has poor branch support in form of a bootstrap value of 50 %. However, when using the UDM-064-LCLR model, the mentioned bootstrap values rise to 100 % (see supplementary data). Also, the C10, C20, to C50 models infer the incorrect topology T1, albeit with decreasing branch support values. Only the maximum likelihood topology of the C60 model is in agreement with the original topology used for simulating the alignment (Figure S28), and has high branch support with bootstrap values of 100 % (see supplementary data).

The improvement in model fit with increasing number of components can also be seen when examining the branch lengths. First, the sum of all branch lengths (*total branch length*) of the original phylogeny used for the simulation is 15.6 average number of substitutions per site. The total branch lengths of the phylogenies estimated by the Poisson, the LG, the UDM-032-LCLR, and the UDM-064-LCLR models are 11.98, 14.14, 14.38, and 14.72 units, respectively. Second, the branch score distance between the original phylogeny used to simulate the alignment and the inferred phylogenies was calculated (Figure S29). The branch score distances of the Poisson and LG model are highest, and the branch score distances of the EDM models decrease with the number of components. For the same number of components the UDM models exhibit lower branch score distances than the CXX models. Ignoring across-site compositional heterogeneity therefore leads to a substantial underestimation of branch lengths because multiple substitution events occurring among a restricted subset of amino-acids are missed by site-homogeneous models.

Also for the simulated data, performance of the UDM models on individual genes was assayed and compared to results of the Poisson and LG models. To this aim, trees were inferred from simulated alignments ranging from 100 to 1000 amino acid columns (see Material and Methods). In general, we observe a decrease of symmetric as well as branch score distance with alignment length (Figures S30, and S31). For short alignments with 100 to 200 columns, an increase of the number of components does not improve the symmetric distance. However, the branch score distance decreases with the number of components, even for such short alignments. For longer alignments, the UDM models with many components consistently outperform the Poisson and LG models, as well as UDM models with fewer components. Strikingly, this trend continues even for UDM models with as many as 1024 components.

## 4 Discussion

The importance of accounting for across-site compositional heterogeneity has been demonstrated by a series of phylogenetic studies where models accounting for across-site compositional heterogeneity, such as the CAT model, were able to overcome artifacts caused by LBA (e.g., Brinkmann et al., 2005; Philippe et al., 2005; Lartillot et al., 2007; Pisani et al., 2015). The reproduction of an LBA artifact and its resolution in a simulation study (Figure 5) provides further evidence for the claim that across-site compositional heterogeneity is a fundamental cause of the phylogenetic artifact observed in the microsporidia data set, and potentially many others.

The simulation study on the microsporidia phylogeny demonstrates that accounting for across-site compositional heterogeneity affects not only the topology (Figures 5 and S28) but also the branch lengths of the inferred phylogeny. In the simulation study, we observe a remarkable downward bias of the total branch length of the phylogeny estimated by the classical Poisson and LG models. The length of long branches, in particular, is severely underestimated. Additionally, the branch score distance between the inferred phylogenies and the original phylogeny used for simulating the alignments improves significantly with the number of EDM model components (Figure S29). Also, we observe superior branch score distances for the UDM models obtained from LCLR transformed site distributions when comparing them to the CXX models. This effect of inadequate modeling of across-site compositional heterogeneity has been overlooked in most previous analyses. With respect to the simulation study, the downward bias of the branch lengths estimated by the Poisson model causes a wrong topology to have higher likelihood than the original topology. This classic LBA attraction artifact is eliminated when accounting for across-site compositional heterogeneity.

In order to provide robust and accurate models that account for across-site compositional heterogeneity, we developed a new method EDCluster to find empirical stationary distributions of amino acids with corresponding weights. EDCluster was used to provide universal stationary distributions estimated from curated databases, but also allows construction of EDM models with a large number of components directly from the data set at hand. The CAT model is employed to infer site distributions, that is, the expectations of the posterior distributions of the stationary distributions of amino acids per site. Subsequently we use a clustering algorithm to explore the structure of the hundred thousands of site distributions. The choice of using a cluster algorithm seemed natural because clustering is a simple machine learning approach for feature discovery. Additionally, to enhance the ability to resolve specialized site distributions we employ coordinate transformations developed specifically for analysis of compositional data. The inference of site distributions with CAT enables our method to deal with the fact that the amino acids of closely related species are expected to be more similar than the ones of distantly related species. Hence, when using our method on an alignment of closely related species the inferred stationary distributions will not necessarily have a low effective number of amino acids. In contrast, methods inferring stationary distributions and weights directly from the alignment (e.g., Susko et al., 2018) require other means to compensate for the expected variation of divergence between the sequences.

From the perspective of potential phylogenetic artifacts caused by inadequate modeling of across-site heterogeneity the effective number of amino acids *K*_eff_ and its distribution provide useful summary statistics for analyzing different models and their stationary distributions. The lower *K*_eff_ is, the higher the potential to underestimate the frequency of multiple substitutions and the probability of homoplasy, with corresponding negative effects on phylogenetic inferences, in terms of recovering accurate branch lengths and avoiding LBA. Consequently, a clustering preceded by a transformation separating stationary distributions with low *K*_eff_, such as the LCLR transformation, can be expected to lead to mixture models less prone to biases in branch length estimation and LBA artifacts. In order to provide sets of universal stationary distributions and weights available for general use we have applied our method, which implements these steps to subsets of databases spanning the whole tree of life. Analysis of the distributions of *K*_eff_ for these universal stationary distributions indicate that a large number of stationary distributions is necessary to adequately model the diversity of site distributions present in empirical alignments. For example, a set of 16 stationary distributions of amino acids is by far not sufficient to describe the observed variety of site distributions (Figure 1). When the number of clusters is too low, we notice that many site distributions are assigned to overly general stationary distributions, because they do not fit in any particular stationary distribution, and not because they are general themselves (Figure 2).

In spite of the apparent need for many stationary distributions, analysis of the WebLogos of the sets of stationary distributions with more than 64 elements reveals an unexpected level of redundancy (see Section S4). It seems reasonable that the number of needed stationary distributions could be reduced by conglomerating stationary distributions exhibiting a certain level of similarity. We attempted to reduce the redundancy within sets comprising many stationary distributions by employing different clustering methods. For example, we tried a form of divisive clustering, where the cluster with the center exhibiting the highest effective number of amino acids is repeatedly divided (Figures S32, and S33), and also density based clustering with DBSCAN (Ester et al., 1996). Both clustering methods failed to improve the redundancy compared to standard *K*-means clustering. However, sets of stationary distributions with a moderate number of elements do not exhibit significant redundancy. For example, the first six elements of the set of 16 stationary distributions obtained from the LCLR transformed site distributions exhibit very little, if any, overlap (Figure 2). Finally, stationary distributions with similar WebLogos may still exhibit specialized features that are not apparent by visual inspection.

A set of stationary distributions and weights together with Poisson exchangeabilities composes an EDM model. We refer to the models composed of the universal stationary distributions and weights discussed above as UDM models. Using the UDM models, we demonstrate the removal of several known LBA associated phylogenetic artifacts from three example analyses: (1) the branching of microsporidia from within fungi and (2) the branching of nematodes and (3) flatworms with arthropodes (Figure 3). For the analysis of the microsporidia data set, the performance of the UDM models was comparable to that of CXX models when using the same number of components. The UDM models outperformed the CXX models in analyses of the data sets including nematodes and platyhelminths. Assaying the ability of different EDM models to adequately recover across-site heterogeneity we found that UDM models outperform CXX models. In fact, the maximum number of components of the CXX models is currently limited to 60 due to the computational cost of the expectation maximization algorithm. In contrast, our method allows for mixture models with many more components. As a proof of concept, we show results for UDM models with 128 and 256 components. All presented analyses support that these component-rich UDM models outperform the C60 model. Another issue with the CXX models is the lack of reproducibility of the expectation maximization estimations in a context characterized by a rugged likelihood surface with a very large number of local maxima (Quang et al., 2008). In particular, the large deviations in the parametric bootstrap results of the C50 model (e.g., Figure 4) reiterate that there may be a problem with respect to local maxima during estimation of the components. The EDCluster approach presented here, however, returns reproducible results, even for rich mixtures.

When examining the total branch lengths of the maximum likelihood phylogenies, we observe that the UDM models obtained from LCLR transformed site distributions exhibit highest total branch lengths. As discussed above, the LCLR transformation facilitates the discovery of more specialized stationary distributions that exhibit lower effective numbers of amino acids. In turn, the lower effective numbers of amino acids lead to inferences exhibiting longer branches. The logarithmic increase of the total branch length with the number of components is striking because it demonstrates that a high number of components may be required. In contrast, the results of the parametric bootstrap analysis indicate that inferences with the UDM-256-LCLR model already overshoot in terms of effective number of amino acids (Figure 4). This observation can be attributed to the LCLR transformation which favors low *K*_eff_ values. In this sense, the LCLR transformation is more eager to catch specific site distributions for moderate numbers of components than the other two transformations but may be too eager for large mixtures of 256 components or more.

The optimal number of components can be determined by established statistical tests, for example, using the AIC or BIC scores. In fact, the maximum log-likelihood as well as the difference in log-likelihood between the tested hypotheses still seem to be far from saturation (Figure 3) for all three transformations. Accordingly, the BIC or AIC scores favor component-rich UDM models, because adding a component only increases the number of model parameters by one (if the weights are inferred). Furthermore, the analyses of the simulated alignments with 100 to 1000 columns show reduced symmetric and branch score distances for component rich UDM models (Figures S30 and S31). These results suggest that the complexity of the composition of site distributions exceeds what can be captured by even the richest mixtures considered here. Consequently, especially for challenging cases, the alleviation of LBA due to site-specific amino-acid preferences may require richer mixtures than the currently available ones such as the CXX models. In summary, for long alignments with thousands of columns, we recommend using UDM models with as many components as permitted by the available computational resources. If the tractable number of components is 256 or lower, we recommend using stationary distributions obtained from the LCLR transformed data, otherwise stationary distributions directly obtained from the data without transformation.

For alignments with 1000 columns or fewer, maximum likelihood estimation of the component weights may be unstable for component rich mixture models, and we recommend Bayesian estimation with UDM models having up to 128 components (Figures S12–S17). If the distribution of component weights is known — for example, from an analysis of the concatenated alignment — maximum likelihood analysis employing component rich UDM models is recommended as it showed high accuracy (Figures S30 and S31).

The results presented above use sets of stationary distributions and weights estimated from a subset of the HOGENOM database. A parallel analysis of a subset of the HSSP database was performed and corresponding components were collected. However, the stationary distributions obtained from the HSSP database were mostly outperformed by the ones obtained from the HOGENOM database. Analysis of the taxonomic composition of the databases (see Materials and Methods) revealed that the taxonomic composition of the subset of the HOGENOM database is enriched for eukaryotes with an approximate value of 70 %, which is in agreement with the taxonomic compositions of the three analyzed data sets. For completeness, the stationary distributions obtained from the HSSP database as well as universal stationary distributions obtained from the union of both databases are also provided, and may exhibit better performance on data sets enriched for bacteria or with a balanced distribution of eukaryotes, archaea, and bacteria, respectively.

We used Poisson exchangeabilities for this first presentation of the UDM models in order to allow comparison with existing models, in particular the CXX models, which were also estimated using Poisson exchangeabilities. In practice, the CXX models are now widely used together with non-uniform exchangeabilities, for example, with the ones of the LG model (a Google scholar search for phylogenetics “LG+C60” returned 59 results on August 22, 2019). However, it may be problematic to use LG exchangeabilities with sets of stationary distributions estimated employing Poisson exchangeabilities, because the effect of across-site compositional heterogeneity might be overfitted. Our method allows estimation of UDM models suitable for a specific set of non-uniform exchangeabilities such as the ones of the LG model by using these exchangeabilities during the inference of the site distributions with the CAT model. Doing so, however, would still raise the question that LG exchangeabilities, originally estimated in a site-homogeneous context, have already captured part of what is in fact a result of site-specific amino-acid preferences. A more principled alternative would be to use the present clustering approach in the context of mutation-selection models (Rodrigue et al., 2010), to estimate universal mixtures of amino-acid fitness profiles.

Although in this contribution we seek to provide a set of models available for universal use, data set specific stationary distributions and weights can be estimated. First, the alignment has to be analyzed with CAT. For this purpose, it is sufficient to fix the topology; a measure greatly reducing computational requirements. If the alignment is still too computationally demanding, it is possibly to split the alignment, or randomly sub-sample a given number of shorter alignments which can be analyzed appropriately (jackknifing). Overall, the computational requirements are much less than a complete analysis with CAT. Second, the site distributions can be analyzed using the provided script (Section S2). Finally, phylogenetic inference can be performed using an EDM model specific to the data set. We tested this procedure on the three discussed data sets and the LBA artifacts were removed in all three cases (see supplementary data).

Before closing the discussion, we would like to examine the relation of EDM models with other available methods. For example, transition rate matrix recoding methods split the amino acids into separate groups representing different physicochemical properties (e.g., Kosiol et al., 2004; Susko et al., 2007). Amino acids within the same group are frequently exchanged whereas there is hardly any exchange between amino acids of different groups. In our opinion, EDM models are very similar in that they differentiate between amino acids exhibiting frequent exchange and amino acids exhibiting no or very limited exchange. However, EDM models seem to be more flexible, because they allow specific amino acids to be member of more than one group, such that the final estimations are superpositions of the individual groupings.

Next, phylogenetic mixture models require a significant amount of computational resources, in particular computer memory. For this reason, the posterior mean site frequency (PMSF, Wang et al., 2018) method has been developed. For each site in the alignment, the PMSF method condenses the stationary distributions of the mixture model components into a single stationary distributions. The single stationary distribution is a weighted superposition of the stationary distributions of all mixture model components. The weights are the posterior probabilities of the site belonging to the respective mixture model components. These posterior probabilities are calculated using a so-called guide tree, which has to be given. The PMSF method speeds up calculations with EDM models and, as such, can be perfectly used with the UDM models.

Similarly, EDM models can be combined with partition models. While, there might be significant evidence justifying the use of specific phylogenetic models for different partitions of the data, this is not in general the case. A canonical way of performing phylogenetic analysis could be: the same EDM model is used across all partitions of the data but a separate set of parameters is inferred for each partition. When using CXX or UDM models, one could only infer a separate set of mixture weights per partition, whereas all other parameters are shared across all partitions.

Finally, usage of an additional mixture model component representing invariable sites (usually +I flag) is possible, but not recommended. First, we did not analyze the effect of this measure. Second, highly constrained stationary distributions with an effective number of amino acids close to 1.0 may already imitate this feature because a very limited availability of amino acids increases the probability of a constant site in the alignment when compared to more general stationary distributions. Additionally, slowly evolving sites are modeled when accounting for across-site rate heterogeneity, for example by a discrete Gamma distribution, which is highly recommended.

Finally, the UDM models can help resolve open phylogenetic problems involving large data sets and distantly related species (e.g., Simion et al., 2017; Philippe et al., 2019). Further, dissimilarities in compositional heterogeneity may be detected by applying EDCluster to specific species groups. Also, the ideal number of EDM model components is still an open question. Statistical tests may not be the best guidance in developing appropriate methods because they favor component-rich EDM models. Albeit, parametric bootstrap analyses, and posterior predictive analyses with Bayesian methods can be used. For EDCluster, automatic clustering algorithms could be used. In conclusion, the presented UDM models constitute a valuable alternative to the widely used CXX models, and can be used for comparisons against the CAT or the CXX models. EDCluster allows estimation of stationary distributions that are specific to the data set at hand and suitable for use with non-uniform exchangeabilities.

## 5 Supplementary material

A supplement to this manuscript is distributed online together with the main text. Supplementary data is available on GitHub at https://github.com/dschrempf/edm-models-data.

## 6 Acknowledgments

D.S. and G.J.Sz. received funding from the European Research Council under the European Union’s Horizon 2020 research and innovation programme under grant agreement no. 714774. DS was also supported by a stipend of excellency of the Austrian government. G.J.Sz. was also supported by the grant GINOP-2.3.2.–15–2016–00057. Some computations were performed on the LBBE/PRABI cluster and on the HPC resources of CINES under the allocation A0040310449 made by GENCI.

## 7 Material and Methods

### Empirical distribution mixture models

Evolution of hereditary characters is assumed to occur according to a mixture of *N* stationary, irreducible, time-continuous Markov processes along a phylogeny 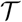. We solely use the state space of amino acids, but the concept of empirical distribution mixture (EDM) models can be applied to arbitrary state spaces of finite cardinality. Let *Q^n^* be the 20×20 transition rate matrix of component *n* with weight *w^n^*. Non-diagonal entries 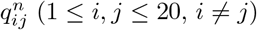 of *Q^n^* can be decomposed into 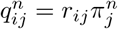. The *r_ij_* are the exchangeabilities which are shared across all components, and *π^n^* is the stationary distribution of component *n*. In this contribution, the Poisson model (Felsenstein, 1981) which exhibits uniform exchangeabilities, was used exclusively. The stationary distributions, which differ between each component, are obtained from curated databases (see below). The diagonal entries 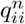 are set such that the row sums are zero. The transition rate matrices are normalized to ensure that one transition of the Markov process is expected to happen per unit length. Additionally, across-site rate heterogeneity can be modeled, for example, by using a discretized Gamma distribution (Yang, 1994a) with parameter *α*. Then, the complete set of EDM model parameters is 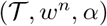. Excluding the phylogeny, EDM models with *N* components have *N* parameters, because ∑_*n*_ *w*^*n*^ = 1.0.

### HOGENOM and HSSP databases

Subsets of the HOGENOM (Dufayard et al., 2005) and HSSP (Schneider et al., 1997) databases consisting of 1005, and 1236 randomly selected alignments were obtained (Quang et al., 2008). For the HOGENOM database, the 1005 alignments contain 15 to 50 sequences and a total number of 175330 amino acid sites. For the HSSP database, the 1236 alignments contain 10 to 100 sequences and a total number of 260961 amino acid sites. Table 1 shows summary statistics of both databases. The summary statistics include the number of sequences and the number sites of the complete databases, and of the analyzed subsets. Further, the percentage of analyzed sites falling into each domain of life provides a rough idea about the taxonomic composition of the analyzed data. First, the analyzed number of sites is slightly larger in the HSSP database compared to the HOGENOM database. Second, and more importantly, the proportion of eukaryotes and bacteria differs largely. The analyzed subset of the HOGENOM database contains a substantially higher proportion of eukaryotes compared to the HSSP database, which comprises a higher proportion of bacteria.

**Table 1:** Size of the HOGENOM and HSSP databases and the analyzed subsets. A rough measure of the taxonomic composition is given in form of the percentage of analyzed sites falling into each domain of life.

|  | **HOGENOM** | **HSSP** |
| --- | --- | --- |
| Number of sequences | 153 818 | 42 999 |
| Number of sites | 40 835 577 | 9 305 643 |
| Number of analyzed sequences | 1005 | 1236 |
| Number of analyzed sites | 175 330 | 260 961 |
| **Distribution across domains in analyzed subsets** |  |  |
| Archaea | 1.6 % | 1.8 % |
| Bacteria | 27.4 % | 59.3 % |
| Eukaryotes | 71.0 % | 35.1 % |
| Viridiae | 0.0 % | 3.8 % |

### Site distributions

For each alignment, a separate Bayesian analysis was conducted using the CAT model (Lartillot et al., 2004) with Poisson exchangeabilities. Phylobayes (Lartillot et al., 2013) was used for the Bayesian analyses. The phylogenies were fixed to the ones estimated by Quang et al. (2008) who had used the WAG model (Whelan et al., 2001) and PhyML (Guindon et al., 2010). Command lines are stated in Section S1. For each alignment and each site, the posterior distribution of the stationary distribution of amino acids is a mapping from the 20-dimensional simplex to the unit interval *p* : *S*^20^ → [0, 1]. The corresponding site distribution, which is the expectation E (*p*), is a point on the 20-dimensional simplex *S*^20^. The site distributions of all sites were collected and used as a basis for all further analysis.

### Transformations of site distributions

The site distributions were analyzed as is, or after transformation from the Aitchison (1982) simplex to real space, which is a standard procedure when analyzing compositional data. First, the well-characterized centered log ratio transformation CLR : *S^d^* → ℝ^*d*^ (Aitchison, 1982) was used. The CLR transformation of a point *x* = (*x*_1_, … , *x*_*d*_) is defined as

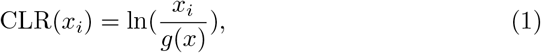

where 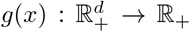 is the geometric mean. Basically, the coordinates of *x* are fanned out from [0, 1]^*d*^ to (−∞, ∞)^*d*^, with the origin (0, …, 0) being CLR ((*g*(*x*), … , *g*(*x*))). Recently, Godichon-Baggioni et al. (2018) reported a novel log centered log ratio transformation LCLR : *S^d^* → ℝ^*d*^ derived from the CLR transformation

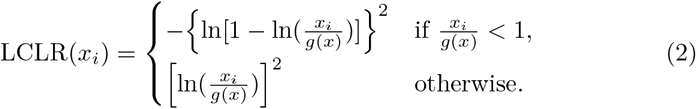

The LCLR transformation moves points that are close to the boundary of the simplex even further away from points that are more in the interior than the CLR transformation. Hence, after the LCLR transformation, points with a low effective number of amino acids (see below) have high Euclidean distances to points with high effective number of amino acids, which is a desired feature.

### Clustering procedure

*K*-means clustering with *K* ∈ {4, 8, 16, …, 256}, was performed on untransformed, CLR-transformed, and LCLR-transformed site distributions with `scikit-learn` (Pedregosa et al., 2011). A maximum number of 500 iterations and a tolerance of 5 × 10*−*5 were used. The stationary distributions of the components of the UDM models are assigned to the obtained cluster centers. The weight of each component is set to the proportion of sites belonging to the respective cluster. During phylogenetic inference, the proposed mixture model weights can be employed without change, or estimated during maximization of the likelihood. In fact, all analyses presented in this manuscript use variable weights estimated during maximization of the likelihood. For details on the EDCluster script used for transforming and clustering the site distributions, please refer to Section S2. In total, we distinguish UDM models with seven different numbers of components, three different types of transformations, and three different databases (HOGENOM, HSSP, and their union). EDCluster, and the obtained stationary distributions and weights are available at https://github.com/dschrempf/edcluster. Sections S3 and S4 present additional analyses of the stationary distributions and weights, and usage instructions for IQ-TREE (Nguyen et al., 2015), Phylobayes, and RevBayes (Höhna et al., 2016), respectively.

### Effective number of amino acids

Metaphorically speaking, entropy is a measure of disorder of a probability distribution. The entropy of a given site distribution *π* is defined as

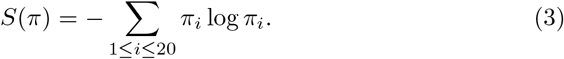

Here, we use the entropy to measure the diversity of a site distribution in the following way

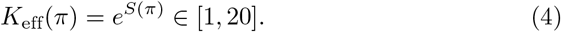

For readability, the explicit dependency on *π* is mostly omitted. We term *K*_eff_ the *effective number of amino acids*, and stationary distributions with high (low) *K*_eff_ *general* (*constrained*). An effective number of amino acids of *K*_eff_ = 1 corresponds to a highly constrained stationary distribution where a single amino acid has probability 1.0, whereas all other amino acids have zero probability. The uniform stationary distribution with *K*_eff_ = 20 is the most general.

### Analyses of data sets

The alignments of the microsporidia, nematode and platyhelminth data sets were obtained from Brinkmann et al. (2005), and from Philippe et al. (2005), respectively. The microsporidia data set contains 40 sequences with a length of 24294 sites. The percentage of gaps is 24.1 % and the average effective number of amino acids is 2.569. The nematode data set contains 37 sequences with a length of 35371 sites. The percentage of gaps is 28.7 % and the average effective number of amino acids is 2.116. The platyhelminth data set contains 32 sequences with a length of 35371 sites. The percentage of gaps is 30.7 % and the average effective number of amino acids is 2.069. The IQ-TREE software package was used for all analyses of the three data sets.

Phylogenetic inference was performed using the WAG, and the LG (Le et al., 2008a) substitution models, the C10 to C60 models (collectively called CXX models; Quang et al., 2008), and the UDM models with 4, 8, 16, *A* …, 256 components. For all analyses, a discrete Gamma distribution with four bins was used to deal with across-site rate heterogeneity (`+G4` model string). For the WAG and LG models, the stationary distribution of amino acids was set to the one observed in the respective alignment. We refrained from adding the stationary distribution of amino acids observed in the data as an additional component to the CXX and UDM models. The weights of the mixture model components of the CXX and UDM models was inferred during maximization of the likelihood. Detailed instruction about how to perform phylogenetic inference with UDM models in IQ-TREE is given in Section S4. For each data set and model, three maximum likelihood analyses were conducted. First, a maximum likelihood analysis inferring the model parameters as well as the topology and the branch lengths of the phylogeny. Further, two analyses with fixed topologies (T1, and T2, see Figure 3) were conducted (`-t` option in IQ-TREE). The branch lengths were inferred without exception.

The analyses of the separated microsporidia genes was performed with Phylobayes. Results for the LG model, the CAT model with Poisson exchangeabilities, and UDM models with four, up to 128 components also with Poisson exchangeabilities were obtained. A burn in of 100 steps, and a total of 1100 steps were used. Rate heterogeneity was accounted for with a discrete gamma distribution with four categories. The inferred trees were compared to two the trees T1, and T2 (Brinkmann et al., 2005). If a gene only contained a subset of all species, the trees T1 and T2 were pruned before comparison such that they contained the same set of species as the respective gene. The symmetric distance (Robinson et al., 1981), the branch score distance Kuhner et al., 1994, and the incompatible split distance were computed. The incompatible split distance is similar to the symmetric distance in that it only accounts for topological differences. However, topological uncertainties do not contribute to the incompatible split distance. Briefly, let us compare two topologies 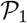 and 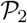, and let *B*_1_ be the only bipartition induced by 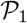 not induced by 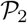. Then, the symmetric distance between 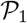 and 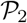 is strictly larger than zero. For the incompatible split distance however, additionally all multifurcations of 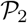 have to be examined. If a multifurcation of 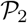 can be resolved such that the resolved order of splits induces *B*_1_, the incompatible split distance is zero.

### Parametric bootstrap analyses

For each data set and phylogenetic model, the maximum likelihood phylogeny and model parameters were used to simulate an alignment with 25 000 sites. For the CXX and UDM models, the stationary distribution at each site is determined randomly from the stationary distributions of the mixture model components using the weights from the respective maximum likelihood inferences. A custom simulator written in Haskell (`elynx`, Section S8) was used for this purpose. Subsequently, the effective number of amino acids *K*_eff_ was calculated per site in the alignment. The obtained distribution of *K*_eff_ values was compared to the one of the original data set using the Wasserstein distance as it is implemented in SciPy (Jones et al., 2001).

### Phylogenetic artifact can be reproduced in simulation study

The phylogeny used to simulate the alignment was chosen from an analysis of the microsporidia data set (Brinkmann et al., 2005) with the UDM model with 64 components obtained from clustering the LCLR transformed site distributions. `elynx` (Section S8) was used to simulate 25 000 sites using Poisson exchangeabilities. Each site was randomly assigned a stationary distribution sampled with replacement from the site distributions of the HOGENOM database which had been obtained by the Bayesian CAT analyses described above. The simulated alignment was analyzed with IQ-TREE using the Poisson model with the empirical distribution observed in the alignment (`Poisson+F` model string), and the UDM models with four up to 64 components obtained from clustering the LCLR transformed site distributions. Ultra fast bootstrap (Hoang et al., 2018) with 1000 samples was used with all models (`−bb 1000` option).

The simulated alignment with 25 000 columns was used to randomly subsample short alignments of length 100, 200, 400, 800, and 1000 columns with replacement. For each length, 25 replicate alignments were sub-sampled. Trees were inferred with IQ-TREE using the Poisson model, the LG model, and UDM models with up to 256 components with fixed component weights. Symmetric and the branch score distances were calculated between the inferred trees and the original tree used for the simulation.

## Notes

#### Summary of Updates

- addition of microsporidia gene analyses - analyses of short simulated alignments - provision of UDM models up to 4096 components - improved train of thought - improved wording

https://github.com/dschrempf/EDCluster

https://github.com/dschrempf/edm-models-data

## References

Aitchison J. 1982. The Statistical Analysis of Compositional Data. J. Royal Stat. Soc. Ser. B (Methological). 44:139–177.

Brinkmann H, Van Der Giezen M, Zhou Y, De Raucourt G P, Philippe H. 2005. An empirical assessment of long-branch attraction artefacts in deep eukaryotic phylogenomics. Syst. Biol. 54:743–757. doi: 10.1080/10635150500234609.

Cavalier-Smith T. 1987. Eukaryotes with no mitochondria. Nature. 326:332–333. doi: 10.1038/326332a0.

Crooks G. E. 2004. WebLogo: A Sequence Logo Generator. Genome Res. 14:1188–1190. doi: 10.1101/gr.849004.

Dufayard J.-F, Duret L, Penel S, Gouy M, Rechenmann F, Perrière G. 2005. Tree pattern matching in phylogenetic trees: automatic search for orthologs or paralogs in homologous gene sequence databases. Bioinformatics. 21:2596–2603. doi: 10.1093/bioinformatics/bti325.

Ester M, Kriegel H.-P, Sander J, Xu X. 1996. A density-based algorithm for discovering clusters in large spatial databases with noise. In: Proceedings of 2nd International Conference on Knowledge Discovery and Data Mining. pp. 226–231.

Felsenstein J. 1981. Evolutionary trees from DNA sequences: a maximum likelihood approach. J. Mol. Evol. 17:368–376.

Felsenstein J. 1978. Cases in which Parsimony or Compatibility Methods will be Positively Misleading. Syst. Biol. 27:401–410. doi: 10.1093/sysbio/27.4.401.

Feuda R, Dohrmann M, Pett W, Philippe H, Rota-Stabelli O, Lartillot N, Wörheide G, Pisani D. 2017. Improved modeling of compositional heterogeneity supports sponges as sister to all other animals. Curr. Biol. 27:3864–3870. doi: 10.1016/j.cub.2017.11.008.

Franzosa E, Xia Y. 2008. Structural Perspectives on Protein Evolution. Annual Reports in Computational Chemistry. Ed. by R A Wheeler, D C Spellmeyer 3–21. doi: 10.1016/s1574-1400(08)00001-7.

Godichon-Baggioni A, Maugis-Rabusseau C, Rau A. 2018. Clustering transformed compositional data using K-means, with applications in gene expression and bicycle sharing system data. J. Appl. Stat. 46:47–65. doi: 10.1080/02664763.2018.1454894.

Goldman N, Thorne J L, Jones D T. 1998. Assessing the impact of secondary structure and solvent accessibility on protein evolution. Genetics. 149:445–58.

Goldman N, Thorne J L, Jones D T. 1996. Using Evolutionary Trees in Protein Secondary Structure Prediction and Other Comparative Sequence Analyses. J. Mol. Biol. 263:196–208. doi: 10.1006/jmbi.1996.0569.

Goldstein R A. 2008. The structure of protein evolution and the evolution of protein structure. Curr. Opin. Struct. Biol. 18:170–177. doi: 10.1016/j.sbi.2008.01.006.

Guindon S, Dufayard J.-F, Lefort V, Anisimova M, Hordijk W, Gascuel O. 2010. New Algorithms and Methods to Estimate Maximum-Likelihood Phylogenies: Assessing the Performance of PhyML 3.0. Syst. Biol. 59:307–321. doi: 10.1093/sysbio/syq010.

Hirt R P, Logsdon J M, Healy B, Dorey M W, Doolittle W F, Embley T M. 1999. Microsporidia are related to Fungi: Evidence from the largest subunit of RNA polymerase II and other proteins. Proc. National Acad. Sci. 96:580–585. doi: 10.1073/pnas.96.2.580.

Hoang D T, Chernomor O, Haeseler A von Minh B Q, Le S V. 2018. UFBoot2: Improving the Ultrafast Bootstrap Approximation. Mol. Biol. Evol. 35:518–522. doi: 10.1093/molbev/msx281.

Höhna S, Landis M J, Heath T A, Boussau B, Lartillot N, Moore B R, Huelsenbeck J P, Ronquist F. 2016. RevBayes: Bayesian phylogenetic inference using graphical models and an interactive model-specification language. Syst. Biol. 65:726–736. doi: 10.1093/sysbio/syw021.

Jimenez M J, Arenas M, Bastolla U. 2018. Substitution Rates Predicted by Stability-Constrained Models of Protein Evolution Are Not Consistent with Empirical Data. Mol. Biol. Evol. 35:743–755. doi: 10.1093/molbev/msx327.

Jones E, Oliphant T, Peterson P, et al. 2001. SciPy: Open source scientific tools for Python.

Jukes T H, Cantor C R. 1969. Evolution of protein molecules. Mammalian Protein Metabolism. Ed. by H N Munro 21–132.

Kamaishi T, Hashimoto T, Nakamura Y, Masuda Y, Nakamura F, Okamoto K.-i, Shimizu M, Hasegawa M. 1996. Complete Nucleotide Sequences of the Genes Encoding Translation Elongation Factors 1 and 2 from a microsporidian parasite, Glugea plecoglossi: Implications for the Deepest Branching of Eukaryotes. J. Biochem. 120:1095–1103. doi: 10.1093/oxfordjournals.jbchem.a021527.

Keeling P J, Fast N M. 2002. Microsporidia: Biology and Evolution of Highly Reduced Intra-cellular Parasites. Annu. Rev. Microbiol. 56:93–116. doi: 10.1146/annurev.micro.56.012302.160854.

Keeling P J, Luker M A, Palmer J D. 2000. Evidence from Beta-Tubulin Phylogeny that Microsporidia Evolved from Within the Fungi. Mol. Biol. Evol. 17:23–31. doi: 10.1093/oxfordjournals.molbev.a026235.

Kosiol C, Goldman N, Buttimore N H. 2004. A new criterion and method for amino acid classification. J. Theor. Biol. 228:97–106. doi: 10.1016/j.jtbi.2003.12.010.

Kuhner M K, Felsenstein J. 1994. A simulation comparison of phylogeny algorithms under equal and unequal evolutionary rates. Mol. Biol. Evol. 11:459–468.

Lanfear R, Frandsen P B, Wright A M, Senfeld T, Calcott B. 2016. PartitionFinder 2: New Methods for Selecting Partitioned Models of Evolution for Molecular and Morphological Phylogenetic Analyses. Mol. Biol. Evol.:msw260. doi: 10.1093/molbev/msw260.

Lartillot N, Brinkmann H, Philippe H. 2007. Suppression of long-branch attraction artefacts in the animal phylogeny using a site-heterogeneous model. BMC Evol. Biol. 7:1–14. doi: 10.1186/1471-2148-7-S1-S4.

Lartillot N, Philippe H. 2004. A Bayesian mixture model for across-site heterogeneities in the amino-acid replacement process. Mol. Biol. Evol. 21:1095–1109. doi: 10.1093/molbev/msh112.

Lartillot N, Rodrigue N, Stubbs D, Richer J. 2013. PhyloBayes MPI: Phylogenetic Reconstruction with Infinite Mixtures of Profiles in a Parallel Environment. Syst. Biol. 62:611–615. doi: 10.1093/sysbio/syt022.

Le S Q, Dang C C, Gascuel O. 2012. Modeling Protein Evolution with Several Amino Acid Replacement Matrices Depending on Site Rates. Mol. Biol. Evol. 29:2921–2936. doi: 10.1093/molbev/mss112.

Le S Q, Gascuel O. 2008a. An improved general amino acid replacement matrix. Mol. Biol. Evol. 25:1307–1320. doi: 10.1093/molbev/msn067.

Le S Q, Gascuel O. 2010. Accounting for Solvent Accessibility and Secondary Structure in Protein Phylogenetics Is Clearly Beneficial. Syst. Biol. 59:277–287. doi: 10.1093/sysbio/syq002.

Le S Q, Lartillot N, Gascuel O. 2008b. Phylogenetic mixture models for proteins. Philos. Trans. Royal Soc. B: Biol. Sci. 363:3965–3976. doi: 10.1098/rstb.2008.0180.

Nguyen L.-T, Schmidt H A, Haeseler A von Minh B Q. 2015. IQ-TREE: A Fast and Effective Stochastic Algorithm for Estimating Maximum-Likelihood Phylogenies. Mol. Biol. Evol. 32:268–274. doi: 10.1093/molbev/msu300.

Pál C, Papp B, Lercher M J. 2006. An integrated view of protein evolution. Nat. Rev. Genet. 7:337–348. doi: 10.1038/nrg1838.

Pedregosa F et al. 2011. Scikit-learn: Machine Learning in Python. J. Mach. Learn. Res. 12:2825–2830.

Philippe H, Brinkmann H, Lavrov D V, Littlewood D T J, Manuel M, Wörheide G, Baurain D. 2011. Resolving Difficult Phylogenetic Questions: Why More Sequences Are Not Enough. PLoS Biol. 9. Ed. by D Penny:e1000602. doi: 10.1371/journal.pbio.1000602.

Philippe H, Lartillot N, Brinkmann H. 2005. Multigene analyses of bilaterian animals corroborate the monophyly of Ecdysozoa, Lophotrochozoa, and protostomia. Mol. Biol. Evol. 22:1246–1253. doi: 10.1093/molbev/msi111.

Philippe H, Laurent J. 1998. How good are deep phylogenetic trees? Curr. Opin. Genet. Dev. 8:616–623. doi: 10.1016/s0959-437x(98)80028-2.

Philippe H et al. 2019. Mitigating Anticipated Effects of Systematic Errors Supports Sister-Group Relationship between Xenacoelomorpha and Ambulacraria. Curr. Biol. 29:1818–1826.e6. doi: 10.1016/j.cub.2019.04.009.

Pisani D, Pett W, Dohrmann M, Feuda R, Rota-Stabelli O, Philippe H, Lartillot N, Wörheide G. 2015. Genomic data do not support comb jellies as the sister group to all other animals. Proc. National Acad. Sci. 112:15402–15407. doi: 10.1073/pnas.1518127112.

Quang L S, Gascuel O, Lartillot N. 2008. Empirical profile mixture models for phylogenetic reconstruction. Bioinformatics. 24:2317–2323. doi: 10.1093/bioinformatics/btn445.

Robinson D F, Foulds L R. 1981. Comparison of phylogenetic trees. Math. Biosci. 53:131–147. doi: 10.1016/0025-5564(81)90043-2.

Rodrigue N, Philippe H, Lartillot N. 2010. Mutation-selection models of coding sequence evolution with site-heterogeneous amino acid fitness profiles. Proc. National Acad. Sci. 107:4629–4634. doi: 10.1073/pnas.0910915107.

Schneider R, Daruvar A de, Sander C. 1997. The HSSP database of protein structure-sequence alignments. Nucleic Acids Res. 25:226–230. doi: 10.1093/nar/25.1.226.

Schwarz G E. 1978. Estimating the Dimension of a Model. The Ann. Stat. 6:461–464. doi: 10.1214/aos/1176344136.

Simion P et al. 2017. A Large and Consistent Phylogenomic Dataset Supports Sponges as the Sister Group to All Other Animals. Curr. Biol. 27:958–967. doi: 10.1016/j.cub.2017.02.031.

Sonnhammer E L, Eddy S R, Durbin R. 1997. Pfam: a comprehensive database of protein domain families based on seed alignments. Proteins. 28:405–20. doi: 10.1002/(SICI)1097-0134(199707)28:3<405::AID-PROT10>3.0.CO;2-L[pii].

Susko E, Lincker L, Roger A J. 2018. Accelerated Estimation of Frequency Classes in Site-Heterogeneous Profile Mixture Models. Mol. Biol. Evol. 35:1–53. doi: 10.1093/molbev/msy026.

Susko E, Roger A J. 2007. On Reduced Amino Acid Alphabets for Phylogenetic Inference. Mol. Biol. Evol. 24:2139–2150. doi: 10.1093/molbev/msm144.

Van de Peer Y, Ben Ali A, Meyer A. 2000. Microsporidia: accumulating molecular evidence that a group of amitochondriate and suspectedly primitive eukaryotes are just curious fungi. Gene. 246:1–8. doi: 10.1016/s0378-1119(00)00063-9.

Vossbrinck C R, Maddox J V, Friedman S, Debrunner-Vossbrinck B A, Woese C R. 1987. Ribosomal RNA sequence suggests microsporidia are extremely ancient eukaryotes. Nature. 326:411–414. doi: 10.1038/326411a0.

Wang H C, Li K, Susko E, Roger A J. 2008. A class frequency mixture model that adjusts for site-specific amino acid frequencies and improves inference of protein phylogeny. BMC Evol. Biol. 8:1–13. doi: 10.1186/1471-2148-8-331.

Wang H C, Minh B Q, Susko E, Roger A J. 2018. Modeling Site Heterogeneity with Posterior Mean Site Frequency Profiles Accelerates Accurate Phylogenomic Estimation. Syst. Biol. 67:216–235. doi: 10.1093/sysbio/syx068.

Whelan N V, Halanych K M. 2016. Who Let the CAT Out of the Bag? Accurately Dealing with Substitutional Heterogeneity in Phylogenomic Analyses. Syst. Biol. 66:232–255. doi: 10.1093/sysbio/syw084.

Whelan S, Goldman N. 2001. A general empirical model of protein evolution derived from multiple protein families using a maximum-likelihood approach. Mol. Biol. Evol. 18:691–699. doi: 10.1093/oxfordjournals.molbev.a003851.

Williams B A P, Hirt R P, Lucocq J M, Embley T M. 2002. A mitochondrial remnant in the microsporidian Trachipleistophora hominis. Nature. 418:865–869. doi: 10.1038/nature00949.

Williams T A, Foster P G, Cox C J, Embley T M. 2013. An archaeal origin of eukaryotes supports only two primary domains of life. Nature. 504:231–236. doi: 10.1038/nature12779.

Yang Z. 1994a. Estimating the pattern of nucleotide substitution. J. Mol. Evol. 39:105–111.

Yang Z. 1994b. Maximum likelihood phylogenetic estimation from DNA sequences with variable rates over sites: approximate methods. J. Mol. Evol. 39:306–314.

