## Supplementary material for "Scalable empirical mixture models that account for across-site compositional heterogeneity"

—

### Supplement

Dominik Schrempf[1], Nicolas Lartillot[2], and Gergely Szöllősi[1,3,4]

[1]Dept. Biological Physics, Eötvös University, Pázmány P. stny. 1A., H-1117 Budapest, Hungary
[2]Laboratoire de Biométrie et Biologie Évolutive, Université Lyon 1, CNRS, UMR, Lyon, France
[3]ELTE-MTA "Lendület" Evolutionary Genomics Research Group, Pázmány P. stny. 1A., H-1117 Budapest, Hungary
[4]Evolutionary Systems Research Group, Centre for Ecological Research, Hungarian Academy of Sciences, H-8237 Tihany, Klebelsberg Kuno str. 3., Hungary

March 31, 2020

**S6 Results for nematodes** **24**

**S7 Results for platyhelminths** **26**

S8 Tool for phylogenetic analysis 27

S9 Parametric bootstrap 28

S10 Simulation study 31

S11 Divisive clustering 36

### S1 Inference of site distributions

Let ALI be a multi-sequence alignment, and TRE be the respective phylogeny which in our case had been inferred using the WAG model (Whelan et al., 2001a). Bayesian inference using Markov chain Monte Carlo was performed with Phylobayes (Lartillot et al., 2013). A burn-in of 100 samples and a total length of the Markov chain of 1100 samples was chosen. The used command line was

```
$ pb_mpi -f -d ALI -T TRE -cat -poisson -x 1 1100 -s ALI_chain
```

Convergence was examined using `tracecomp`. For example, the output for the first alignment `A00001` of the HOGENOM data base (Dufayard et al., 2005) obtained from Quang et al. (2008) was

```
$ tracecomp -x 100 A00001_chain{1,2}.trace
setting upper limit to : 1100
A00001_chain1.trace  burnin : 100 sample size : 1000
A00001_chain2.trace  burnin : 100 sample size : 1000
name                 effsize rel_diff

loglik               51     0.527818
length               166    0.147459
alpha                67     0.543103
pinv                 1000   0
Nmode                122    0.269335
statent              36     0.600935
statalpha            86     0.361928
kappa                329    0.242377
```

The expectations of the posterior distribution of the stationary distribution per site, which are called site distributions (see main text), were obtained using `readpb_mpi`

```
$ readpb_mpi -ss -x 100 1 ALI_chain
```

### S2 Clustering of site distributions

The site distributions were transformed and clustered using EDCluster which is provided at https://github.com/dschrempf/edcluster.
EDCluster uses `scikit-learn` (Pedregosa et al., 2011). Of course, EDCluster can also be applied to site distributions obtained from a specific data set. For further instructions, please refer to the documentation delivered together with the script. The usage is as follows.

Help of clustering script

```
usage: EDCluster [-h] [-t {none,clr,lclr}] [-k K] [-n] [-p P] [--pickle-only] [--continue-run] [-v] [filenames [filenames ...]]

EDCluster version 2.0.0.
Developed by Dominik Schrempf (2020).

Scalable detection of empirical distribution mixture models by
transformation and clustering of site distributions. Each amino acid frequency
distribution is considered as a point in twenty dimensional space. The K-Means
clustering algorithm is used to label the points from 1 to K, for a given value
of K. The clusters are output in various file formats, readily usable with
IQ-TREE [1], Phylobayes [2], and RevBayes [3].

Before clustering, coordinate transformations can be applied.

Implemented transformations:
- centered log ratio (CLR) transform [4]
- log centered log ratio (LCLR) transform [5]

Points close to the boundaries, that is, with some coordinates having low
probability, exhibit special features. The coordinate transformations project
these points to a position far away from the projection of points with high
diversity and intermediate frequency values for all coordinates.

The site distribution files are expected to be tab separated, and of the
following form:

----------------------------------------------------------------------
  A C D E F G H I K L M N P Q R S T V W Y

1 0.008 0.001 0.001 0.007 0.002 0.004 0.01 ...
2 ...
...
----------------------------------------------------------------------

By default, EDcluster calculates mixture model components of empirical
distribution mixture (EDM) models for K={4,8,16,32,64,128,256,512,1024,2048},
and all coordinate transformations. This can take a while. Calculation of an
EDM model with a specific number of components, as well as a specific
transformation can be achieved with the -k, and -t options (see below).

positional arguments:
  filenames           Names of files containing site distributions (default: None)

optional arguments:
  -h, --help          show this help message and exit
  -t {none,clr,lclr}  Use specific transformation. (default: None)
  -k K                Use specific number of clusters. (default: None)
  -n                  Also create Nicologos. (default: False)
  -p P                Prefix of output files. (default: None)
  --pickle-only       Read, transform, and save data; then exit. (default: False)
```

```
  --continue-run      Use saved data for clustering. (default: False)
  -v                  show program's version number and exit

[1] Nguyen, L., Schmidt, H. A., von Haeseler, A., & Minh, B. Q. (2015).
Iq-tree: a fast and effective stochastic algorithm for estimating
maximum-likelihood phylogenies. Molecular Biology and Evolution, 32(1),
268–274. http://dx.doi.org/10.1093/molbev/msu300

[2] Lartillot, N., Rodrigue, N., Stubbs, D., & Richer, J. (2013). Phylobayes
mpi: phylogenetic reconstruction with infinite mixtures of profiles in a
parallel environment. Systematic Biology, 62(4), 611–615.
http://dx.doi.org/10.1093/sysbio/syt022

[3] Höhna, S., Landis, M. J., Heath, T. A., Boussau, B., Lartillot, N., Moore,
B. R., Huelsenbeck, J. P., … (2016). Revbayes: bayesian phylogenetic inference
using graphical models and an interactive model-specification language.
Systematic Biology, 65(4), 726–736. http://dx.doi.org/10.1093/sysbio/syw021

[4] Aitchison, J. (1982). The statistical analysis of compositional data.
Journal of the Royal Statistical Society, Series B (Methological), 44(2),
139–177.

[5] Godichon-Baggioni, A., Maugis-Rabusseau, C., & Rau, A. (2017).
Clustering transformed compositional data using K-means, with applications in
gene expression and bicycle sharing system data. ArXiv, 1–32.
```

### S3 Universal distribution mixture model components

This section provides further results of the $K$-means clustering of the untransformed, center log ratio (CLR; Aitchison, 1982) transformed, and log center log ratio (LCLR; Godichon-Baggioni et al., 2018) transformed site distributions. The obtained universal distribution mixture (UDM) models can be downloaded from `https://github.com/dschrempf/edcluster`. For instruction on how to use the stationary distributions, see Section S4.

In general, we observe that the more stationary distributions are used, the lower is the average effective number of amino acids (see Materials and Methods in the main text). Next to the measurement using entropy, we can also measure the effective number of amino acids using homoplasy. Homoplasy, in the narrower sense of phylogenetics, is when characters observed at a specific site in two separate and significantly diverged species are equal. For a given site distribution $\pi$, the probability of homoplasy is just the probability of random equality of characters. For amino acids, we have

$$G(\pi) = \sum_{1 \leq i \leq 20} \pi_i^2. \tag{S1}$$

The probability of homoplasy can also be used to measure the diversity of a site distribution

$$K_{\text{eff}}^{H}(\pi) = G(\pi)^{-1} \in [1, 20]. \tag{S2}$$

#### S3.1 No transformation

The effective number of amino acids of the stationary distributions obtained from site distributions without coordinate transformation tend to be higher than the ones with applied coordinate transformations. The trend “more stationary distributions, lower effective number of amino acids” can already be observed well (Figure S1). Also, the total number of required stationary distributions and weights (a stationary distribution with corresponding weight is referred to as *component*) to capture the distribution of site distributions obtained from the CAT analysis of the data seems to be high. Especially, see the notable difference between 64 and 256 components (Figure S2).

No transformation

```
Number of clusters: 4
Index  Weight  Keff_entropy  Keff_homoplasy
0       0.448        17.541          15.941
1       0.302         8.051           5.635
2       0.204        10.871           7.292
3       0.046         3.984           2.048
Weighted average of Keff_entropy across indices: 12.70
Weighted average of Keff_homoplasy across indices: 10.43
Number of clusters: 8
Index   Weight  Keff_entropy  Keff_homoplasy
0       0.306        16.976          14.528
1       0.154         6.879           4.587
2       0.133         7.129           4.227
3       0.104         9.564           6.167
4       0.101        10.208           6.348
5       0.088         7.686           4.490
6       0.069         7.945           4.790
7       0.045         3.945           2.028
Weighted average of Keff_entropy across indices: 10.63
Weighted average of Keff_homoplasy across indices: 7.81
Number of clusters: 16
Index  Weight  Keff_entropy  Keff_homoplasy
0       0.156        17.062          15.521
1       0.107         9.062           6.191
2       0.102         5.493           3.702
3       0.084        14.288          11.434
4       0.076         8.795           5.242
5       0.069        10.448           6.427
6       0.064         7.466           4.482
7       0.062         5.077           2.864
8       0.060         6.665           3.726
9       0.058         8.828           5.269
10      0.044         3.887           2.005
11      0.032         7.244           3.532
12      0.029         6.933           3.763
13      0.027        11.183           6.921
14      0.025         7.943           4.472
15      0.006         6.107           2.879
Weighted average of Keff_entropy across indices: 9.62
Weighted average of Keff_homoplasy across indices: 6.88
```

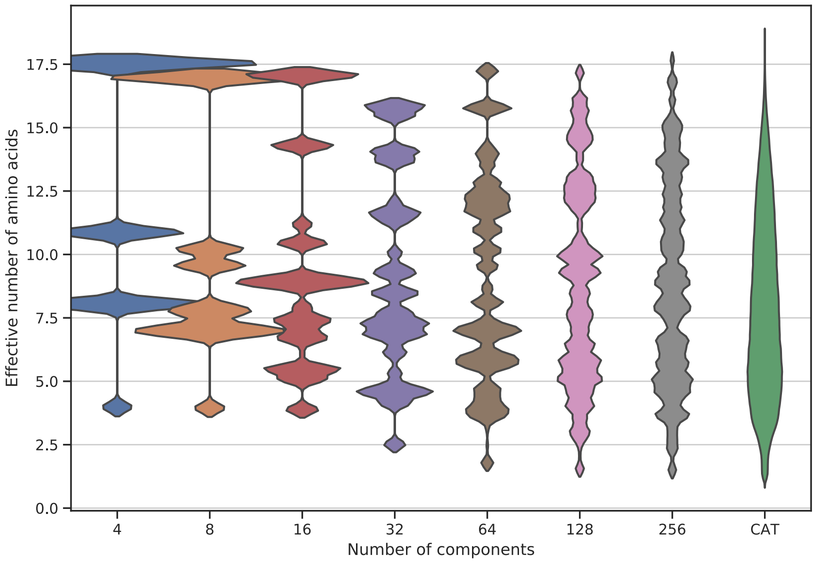

**Figure S1:** Violin plot of the distribution of the effective number of amino acids of the stationary distributions and weights (components) obtained from original site distributions (Godichon-Baggioni et al., 2018) of the HOGENOM database (Dufayard et al., 2005) for a different number of components. On the far right, the distribution of the effective number of amino acids of the site distributions obtained with the CAT model (Lartillot et al., 2004) using Poisson exchangeabilities (Felsenstein, 1981) is shown. The width of the violin plots was normalized such that all areas are equal.

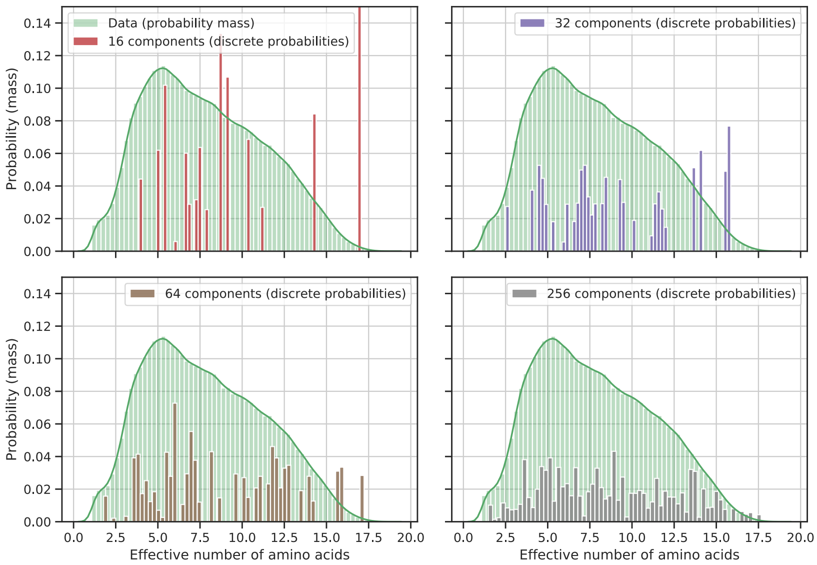

**Figure S2:** Distribution of the effective number of amino acids of the site distributions inferred by the CAT model (Lartillot et al., 2004) using Poisson exchangeabilities (Felsenstein, 1981) from the HOGENOM database (Dufayard et al., 2005), and of the stationary distributions and weights (components) for a different numbers of components obtained from clustering the site distributions without transformation. More components capture the distribution of the site distributions in a better way.

#### S3.2 Center log ratio transformation

The results of the clustering of the CLR transformed site distributions seem to improve upon the results without transformation. In particular, the effective number of amino acids is lower (see the following listing, and Figure S3), and the distribution of effective number of amino acids of the site distribution obtained from the HOGENOM database is modeled in a better way (Figure S4). Nevertheless, it seems that many components are necessary.

CLR transformation

```
Number of clusters: 4
Index  Weight  Keff_entropy  Keff_homoplasy
0       0.369        14.815          12.929
1       0.280         6.621           4.836
2       0.237         9.895           6.533
3       0.114        10.524           6.745
Weighted average of Keff_entropy across indices: 10.87
Weighted average of Keff_homoplasy across indices: 8.44
Number of clusters: 8
Index  Weight  Keff_entropy  Keff_homoplasy
0       0.211         5.984           4.472
1       0.157        11.135           8.096
2       0.148        12.092           8.425
3       0.118        13.456           9.999
4       0.107        11.729           8.582
5       0.105         7.405           4.759
6       0.092         8.484           5.146
7       0.062         4.509           2.363
Weighted average of Keff_entropy across indices: 9.48
```

```
Weighted average of Keff_homoplasy across indices: 6.68
Number of clusters: 16
Index  Weight  Keff_entropy  Keff_homoplasy
0       0.088         5.233           3.334
1       0.084         4.468           3.233
2       0.079         8.327           5.004
3       0.076         9.815           6.177
4       0.076        13.490          11.353
5       0.072         7.716           4.836
6       0.065        13.394          10.378
7       0.061        12.001           9.052
8       0.055         5.227           3.284
9       0.055         9.252           5.168
10      0.055         9.885           7.382
11      0.052         6.243           3.760
12      0.051        17.086          15.103
13      0.044         8.716           5.931
14      0.044         3.102           1.731
15      0.043         7.063           5.426
Weighted average of Keff_entropy across indices: 8.79
Weighted average of Keff_homoplasy across indices: 6.28
```

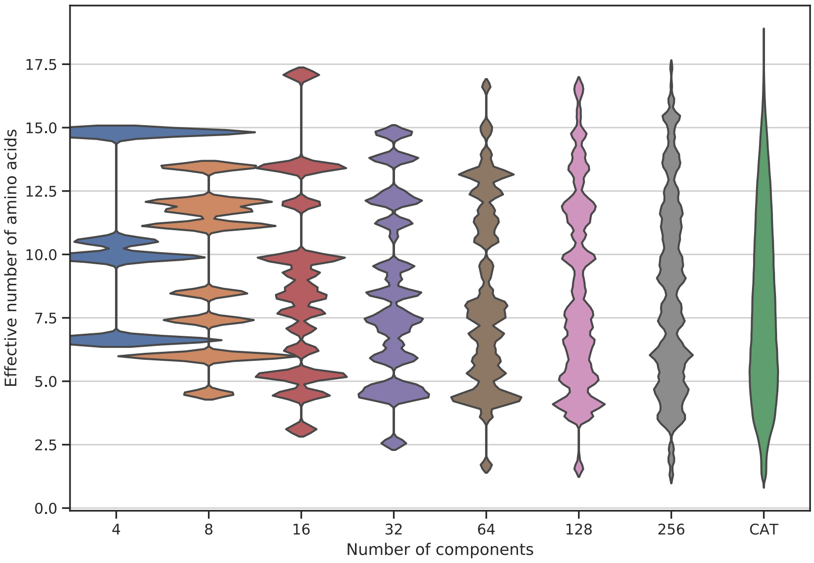

**Figure S3:** See Figure S1 but with center log ratio transformation (Aitchison, 1982).

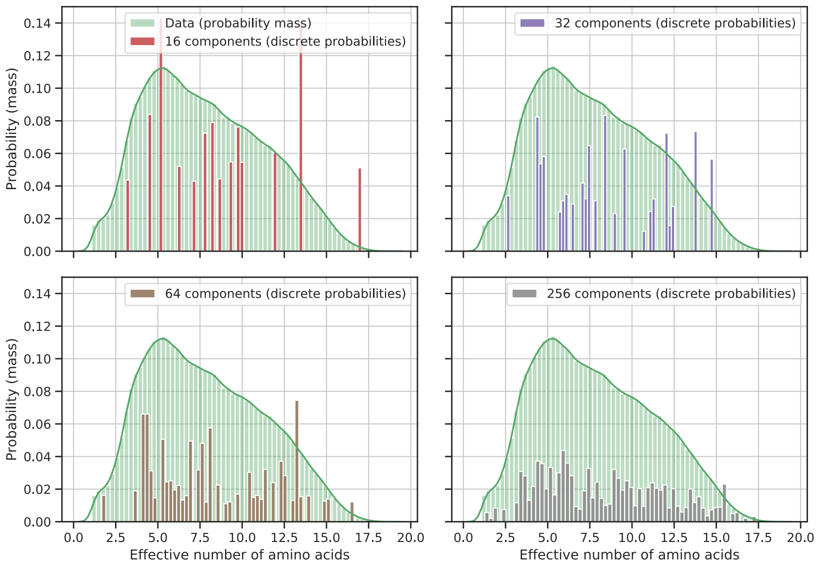

**Figure S4:** See Figure S2 but with center log ratio transformation (Aitchison, 1982).

#### S3.3 Log center log ratio transformation

Most of the results of the clustering of site distributions obtained from the LCLR transformed site distributions are presented in the main text. We observe, that even for 16 components, only one component has a high effective number of amino acids (the general component).

LCLR transformation

```
Number of clusters: 4
Index  Weight  Keff_entropy  Keff_homoplasy
0       0.595        16.394          15.053
1       0.229         6.133           4.613
2       0.102         6.120           4.175
3       0.074         7.282           4.609
Weighted average of Keff_entropy across indices: 12.32
Weighted average of Keff_homoplasy across indices: 10.78
Number of clusters: 8
Index  Weight  Keff_entropy  Keff_homoplasy
0       0.395        17.285          15.892
1       0.137         6.148           3.982
2       0.101         4.768           3.444
3       0.101         7.004           4.626
4       0.090         8.484           5.564
5       0.082         8.469           5.291
6       0.058         6.267           3.920
7       0.036         2.417           1.480
Weighted average of Keff_entropy across indices: 10.77
Weighted average of Keff_homoplasy across indices: 8.86
Number of clusters: 16
Index  Weight  Keff_entropy  Keff_homoplasy
0       0.272        17.064          15.810
1       0.083        10.051           6.866
2       0.082         8.923           5.991
3       0.070         7.800           4.811
4       0.067         7.313           5.016
```

```
5       0.061         7.952          4.913
6       0.051         5.821          3.657
7       0.050         5.436          4.162
8       0.048         4.246          2.563
9       0.043         3.721          2.794
10      0.035         4.354          2.732
11      0.033         5.200          2.653
12      0.031         6.387          3.971
13      0.030         6.083          3.028
14      0.028         6.612          3.847
15      0.015         1.586          1.186
Weighted average of Keff_entropy across indices: 9.58
Weighted average of Keff_homoplasy across indices: 7.50
```

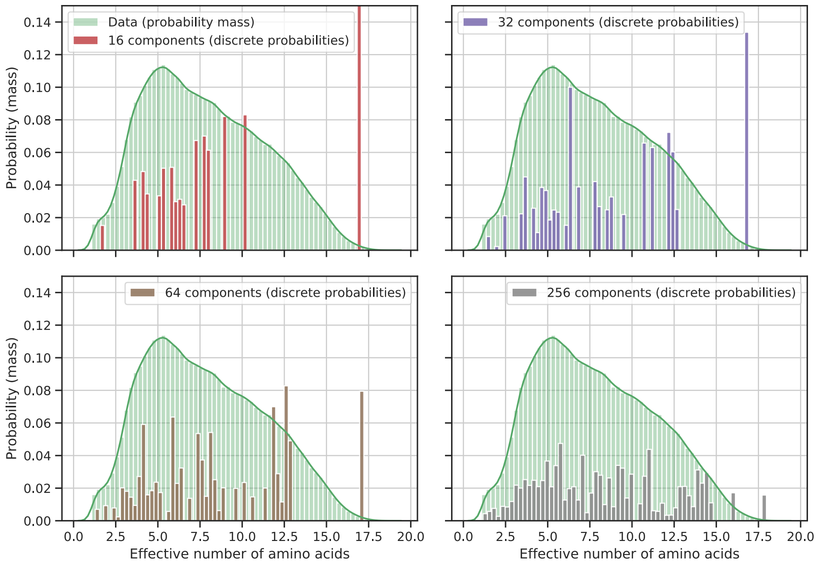

**Figure S5:** See Figure S2 but with log center log ratio transformation (Godichon-Baggioni et al., 2018).

The first, and most general component is a "catch all" case. Many data points are associated with it, because there is no stationary distribution which is more suitable, that is, closer when measured in Euclidian distance in the transformed space. However, many site distributions are not assigned to the general component because they are particularly close (Figure S6-S8). Consequently, the distribution of effective number of amino acids of the site distributions associated with the first component is biased to lower values.

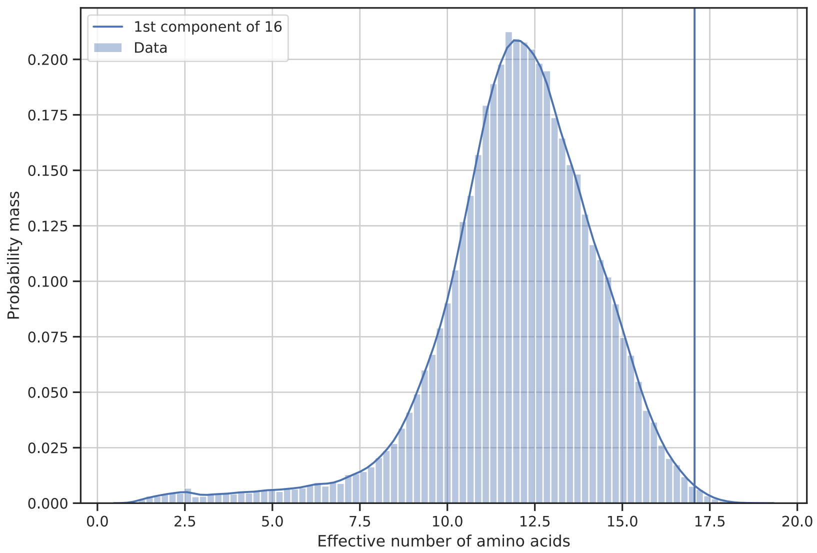

**Figure S6:** Kerndel density estimation of the distribution of effective number of amino acids for site distributions associated with the first component of a total of 16 components, and the effective number of amino acids of the first component itself are shown.

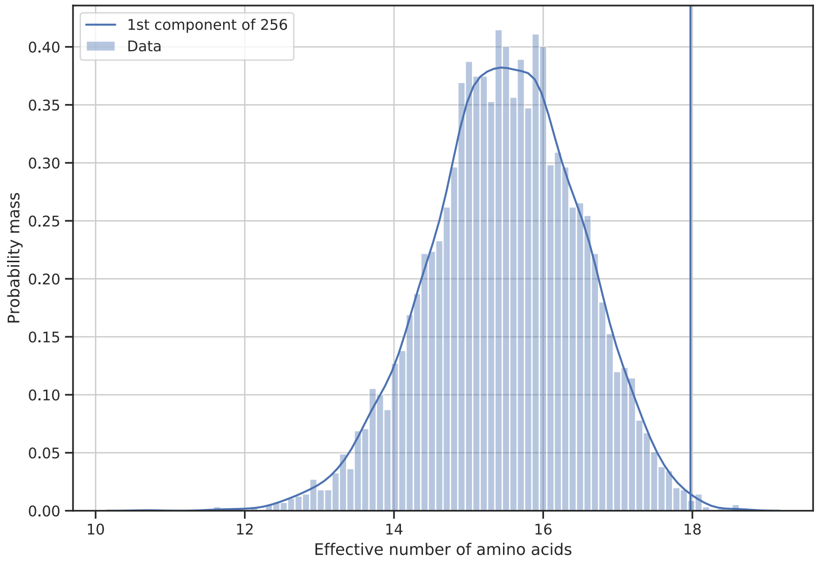

**Figure S7:** See Figure S6, but with 256 components.

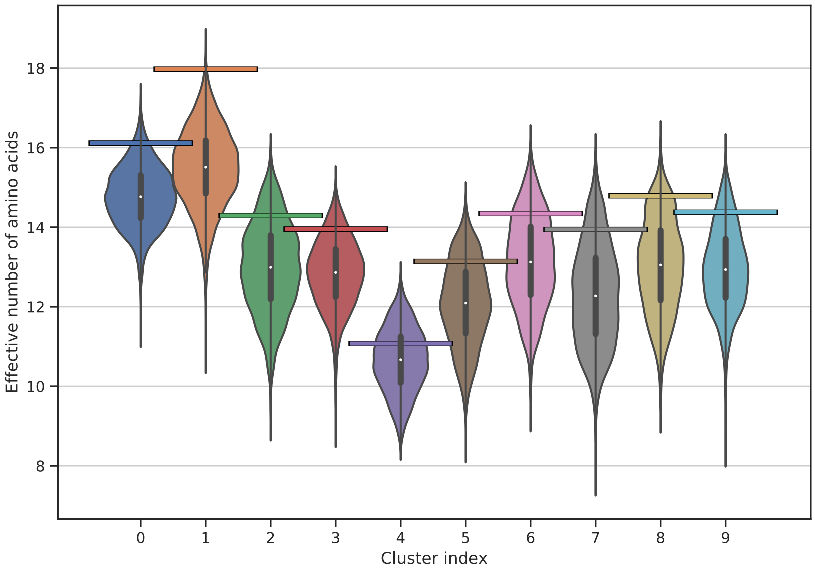

**Figure S8:** Violin plot of the effective number of amino acids of the site distributions associated with the first ten components sorted by weight of the universal distribution mixture model with a total number of 256 components obtained from clustering the log center log ratio (Godichon-Baggioni et al., 2018) transformed site distributions. The actual effective number of amino acids of the stationary distribution of each component is shown by a horizontal line.

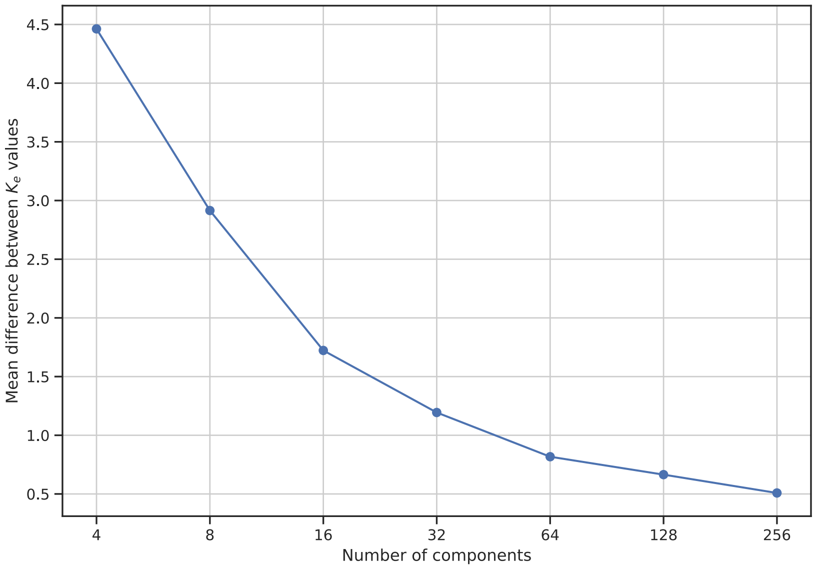

**Figure S9:** Mean difference of the effective number of amino acids of the site distributions and their associated cluster centers for universal distribution mixture models with a different number of components obtained from clustering the log center log ratio (Godichon-Baggioni et al., 2018) transformed site distributions. Values greater than zero imply that the $K_{\text{eff}}$ values of the cluster centers are, on average, more general than the ones of the site distributions.

### S4 Usage of stationary distributions

The files are named in the following format:

```
udm_${database}_${nclusters}_${transformation}_${software}.extension
```

For each combination of parameters (data base, number of parameters, transformation), three files are provided; one for each of the following software packages

- IQ-TREE (Nguyen et al., 2015),
- Phylobayes (Lartillot et al., 2013), and
- RevBayes (Höhna et al., 2016).

Additionally, postscript files show customized WebLogos (Crooks, 2004) of the distributions (see main text). For example, the components as well as the corresponding logos of the UDM model with four components obtained from the HOGENOM database with the LCLR transformation are listed in these files:

```
udm_hogenom_0004_lclr_iqtree.nex
udm_hogenom_0004_lclr_pb.txt
udm_hogenom_0004_lclr_rb.txt
udm_hogenom_0004_lclr_nicologo.ps
```

Additionally, for IQ-TREE all distributions are conveniently summarized into a file `udm_models_${transformation}.nex`. The following command line examples show how these files can be used. The analyzed multisequence alignment is again abbreviated by `ALI`.

#### S4.1 IQ-TREE

The `*iqtree.nex` file defines a frequency mixture model (FMIX). The FMIX model is called, for example, `UDM0004LCLR`. The corresponding analysis using the UDM-4-LCLR model can be run with

```
iqtree -s ALI -mdef udm_models_hogenom.nex -m Poisson+UDM0004LCLR -mwopt
```

Of course, other exchangeabilities such as the ones of the LG model can be used, although we do not advice the usage of non-uniform exchangeabilities, and did not test the performance of non-uniform exchangeabilities extensively.

#### S4.2 Phylobayes

The `*pb.txt` file defines the components. A UDM model analysis can, for example, be run with

```
pb_mpi -d ALI -poisson -catfix udm_hogenom_0004_lclr_pb.txt -s CHAIN_NAME
```

#### S4.3 RevBayes

The `*rb.txt` defines variables to be used in the RevBayes analysis:

**`distributions`** Array containing stationary distributions.

**`distribution_weights`** Array containing weights.

**`n_clusters`** Integer with number of clusters.

A sample script using the UDM model with exchangeabilities of the general time reverible model (GTR; Tavaré, 1986) and four components obtained from the HOGENOM database using the LCLR transformation is given below.

```
data       = readDiscreteCharacterData(ALI)

taxa       <- data.taxa()
n_species  <- data.ntaxa()
n_branches <- 2 * n_species - 3

moves      = VectorMoves()
monitors   = VectorMonitors()

# UDM model.
source("udm_hogenom_0004_lclr_rb.txt")

# Exchangeabilities and moves.
er_prior_alpha <- rep(1, 190)
er ~ dnDirichlet(er_prior_alpha)
moves.append(mvBetaSimplex(er, weight=190))
moves.append(mvDirichletSimplex(er, weight=19))

# Substitution rate matrices and moves.
for ( i in 1:n_clusters ) {
   Q[i] := fnGTR(er, simplex(distributions[i]) )
```

```
}
weights ~ dnDirichlet(rep(1, n_clusters ))
moves.append(mvSimplexElementScale(weights, alpha=10, weight=2.0))

# Prior distribution on the tree topology.
topology ~ dnUniformTopology(taxa)
moves.append(mvNNI(topology, weight=n_species))
moves.append(mvSPR(topology, weight=n_species/10.0))

# Branch length prior.
for (i in 1:n_branches) {
    bl[i] ~ dnExponential(10.0)
    moves.append(mvScale(bl[i]))
}

# Tree.
psi := treeAssembly(topology, bl)

# Connect substitution and tree models.
seq ~ dnPhyloCTMC(tree=psi, Q=Q, type="AA", siteMatrices=weights)
# Attach data.
seq.clamp(data)

# Analysis.
mymodel = model(psi)

# Monitors.
fn_out_base = "output/" + fn_prefix + "_" + fn_distributions
monitors.append(mnFile(psi, filename=fn_out_base + ".trees", printgen=1))
monitors.append(mnModel(filename=fn_out_base + ".log", printgen=1))

# Run.
mymcmc = mcmc(mymodel, moves, monitors)
mymcmc.burnin(generations=1000, tuningInterval=200)
mymcmc.run(generations=3000)

q()
```

### S5 Results for microsporidia

#### S5.1 Concatenated alignments

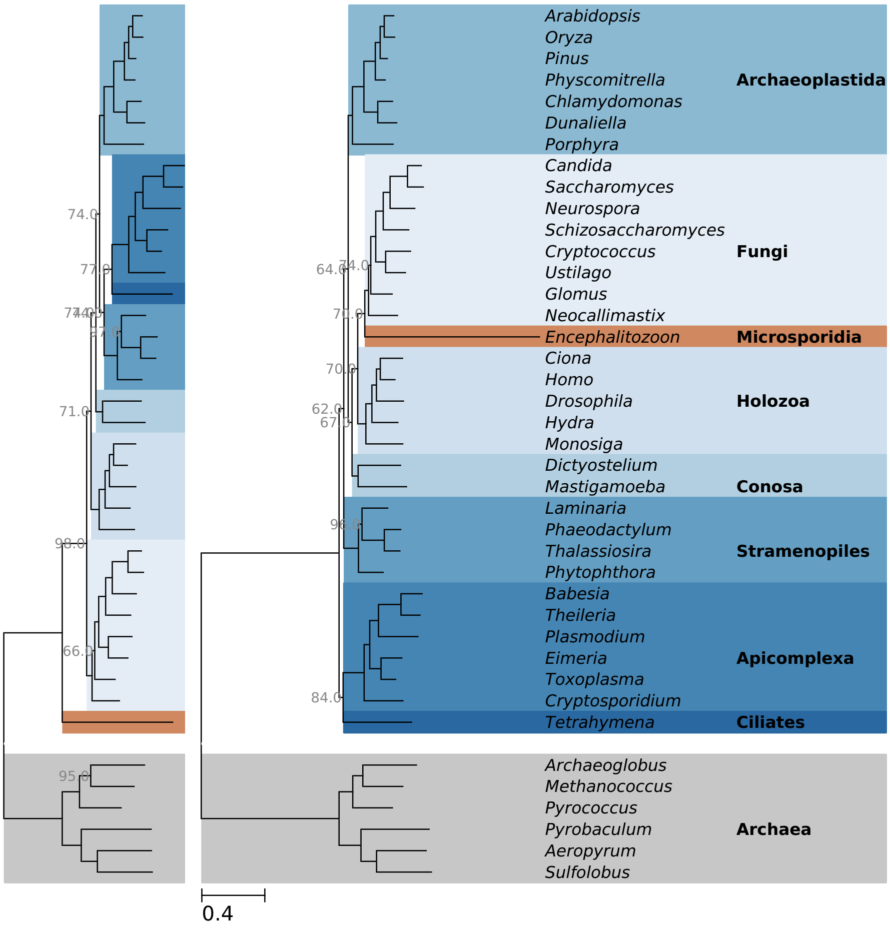

**Figure S10:** Inferred maximum likelihood phylogenies for the microsporidia data set (Brinkmann et al., 2005). (Left) LG model (Le et al., 2008). (Right) Universal distribution mixture model with 64 components obtained from the log center log ratio (Godichon-Baggioni et al., 2018) transformed site distributions. For both phylogenies, branch support estimated by bootstrapping is shown for not fully supported branches.

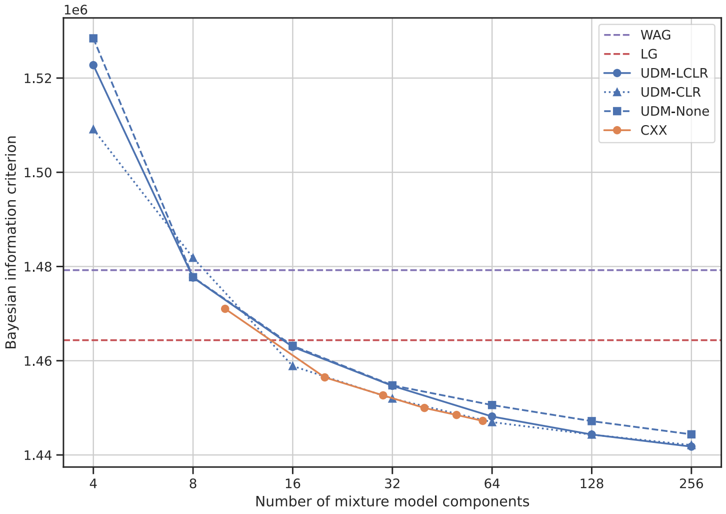

**Figure S11:** Bayesian information criterion (BIC; Schwarz, 1978) scores for different empirical distribution mixture models, the WAG model (Whelan et al., 2001b), and the LG model (Le et al., 2008) applied to the microsporidia data set (Brinkmann et al., 2005).

#### S5.2 Separate genes — comparison to historical tree T1

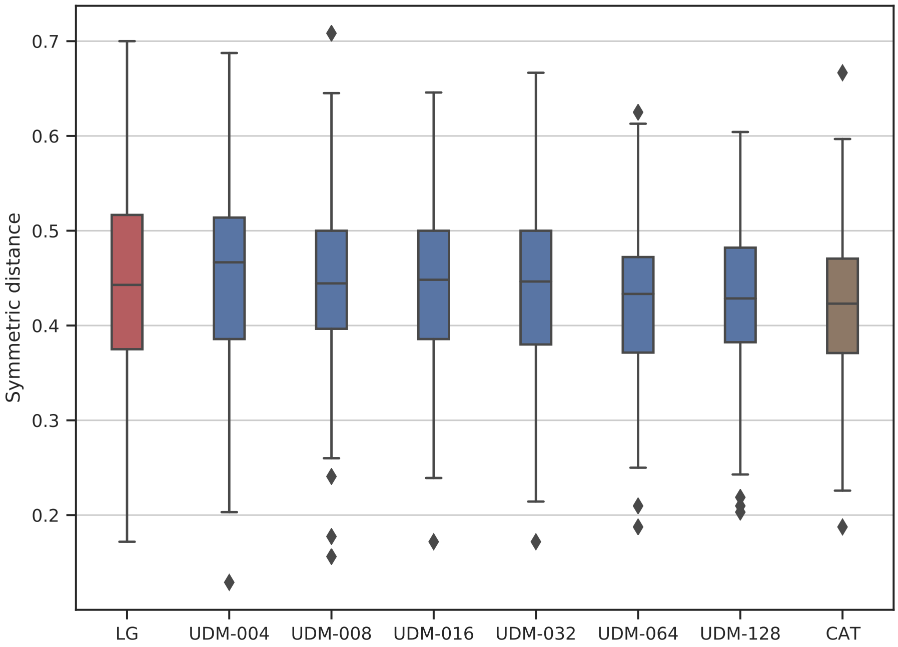

**Figure S12:** Normalized symmetric distances (Robinson et al., 1981) obtained from the gene alignments of the microsporidia data set (Brinkmann et al., 2005) when compared to the historical tree T1 (Figure 3). Results for the LG model (Le et al., 2008), universal distribution mixture models with four to 128 components obtained from the log center log ratio (Godichon-Baggioni et al., 2018) transformed site distributions, and the CAT model (Lartillot et al., 2004) with Poisson exchangeabilities are shown.

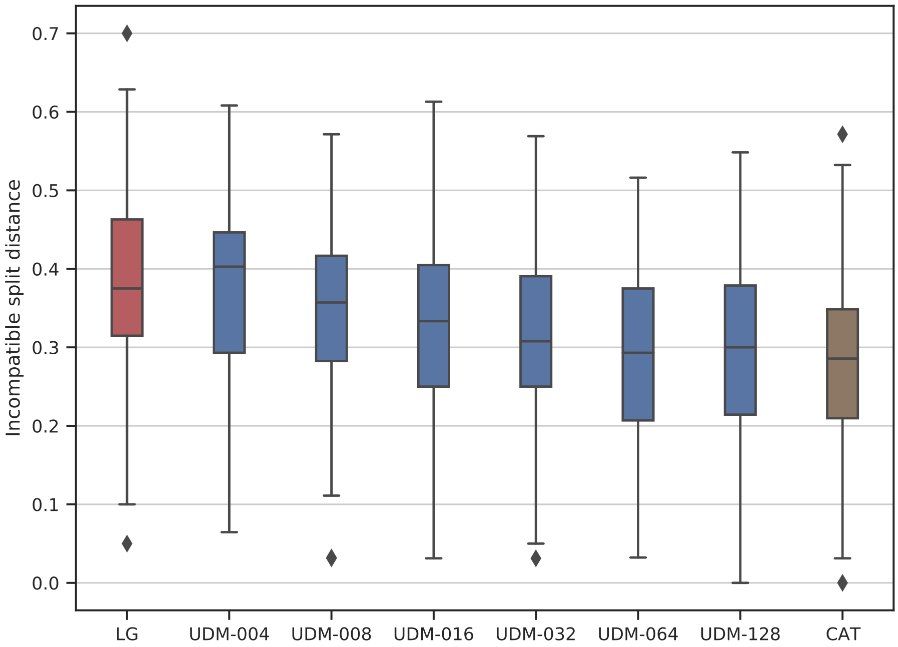

**Figure S13:** Normalized incompatible split distances (see Material and Methods in the main text) obtained from the gene alignments of the microsporidia data set (Brinkmann et al., 2005) when compared to the historical tree T1 (Figure 3). Results for the LG model (Le et al., 2008), universal distribution mixture models with four to 128 components obtained from the log center log ratio (Godichon-Baggioni et al., 2018) transformed site distributions, and the CAT model (Lartillot et al., 2004) with Poisson exchangeabilities are shown.

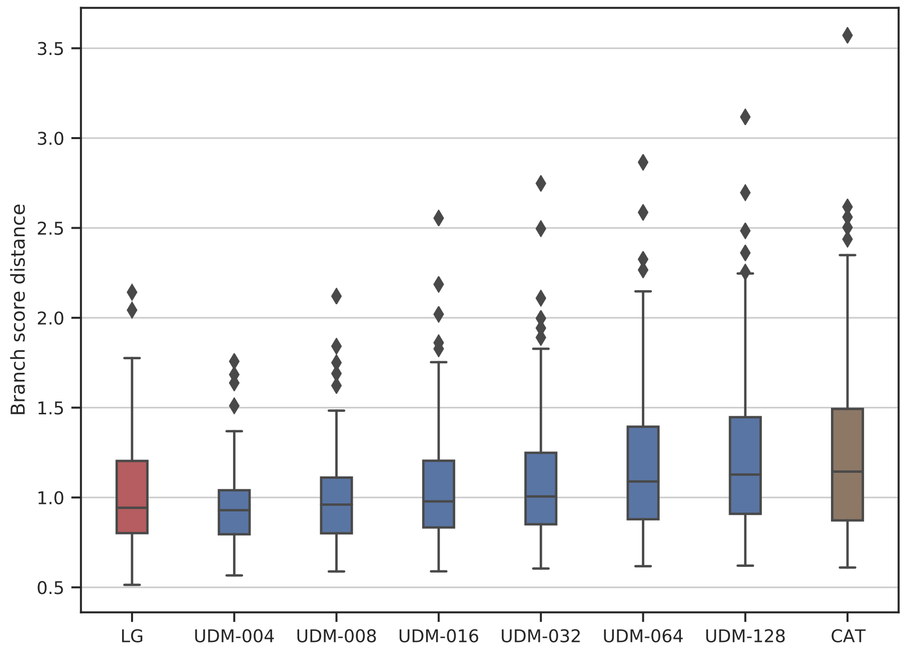

**Figure S14:** Branch score distances (Kuhner et al., 1994) obtained from the gene alignments of the microsporidia data set (Brinkmann et al., 2005) when compared to the historical tree T1 (Figure 3). Results for the LG model (Le et al., 2008), universal distribution mixture models with four to 128 components obtained from the log center log ratio (Godichon-Baggioni et al., 2018) transformed site distributions, and the CAT model (Lartillot et al., 2004) with Poisson exchangeabilities are shown.

#### S5.3 Separate genes — comparison to currently accepted tree T2

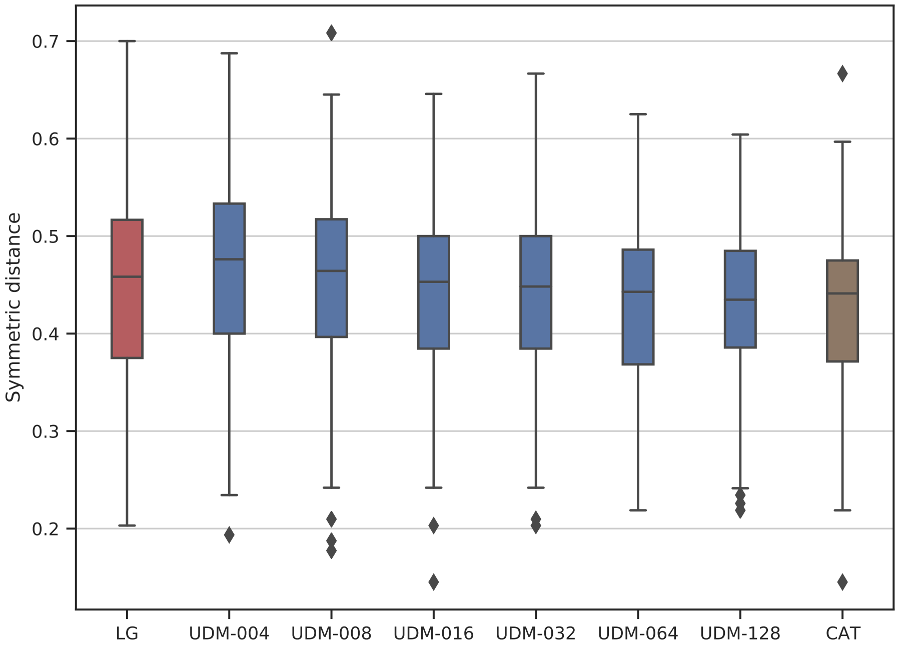

**Figure S15:** Normalized symmetric distances (Robinson et al., 1981) obtained from the gene alignments of the microsporidia data set (Brinkmann et al., 2005) when compared to the currently accepted tree T2 (Figure 3). Results for the LG model (Le et al., 2008), universal distribution mixture models with four to 128 components obtained from the log center log ratio (Godichon-Baggioni et al., 2018) transformed site distributions, and the CAT model (Lartillot et al., 2004) with Poisson exchangeabilities are shown.

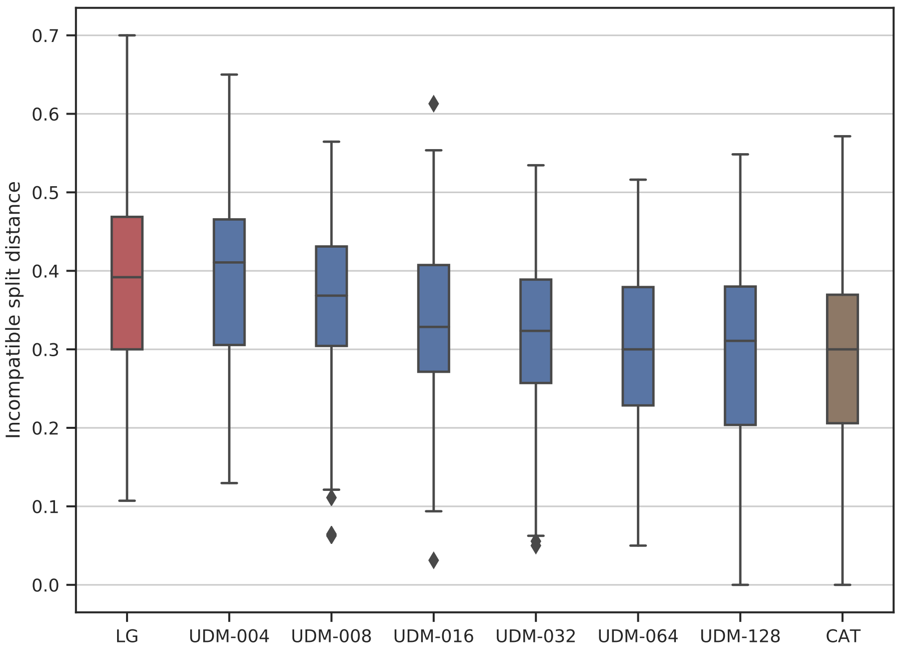

**Figure S16:** Normalized incompatible split distances (see Material and Methods in the main text) obtained from the gene alignments of the microsporidia data set (Brinkmann et al., 2005) when compared to the currently accepted tree T2 (Figure 3). Results for the LG model (Le et al., 2008), universal distribution mixture models with four to 128 components obtained from the log center log ratio (Godichon-Baggioni et al., 2018) transformed site distributions, and the CAT model (Lartillot et al., 2004) with Poisson exchangeabilities are shown.

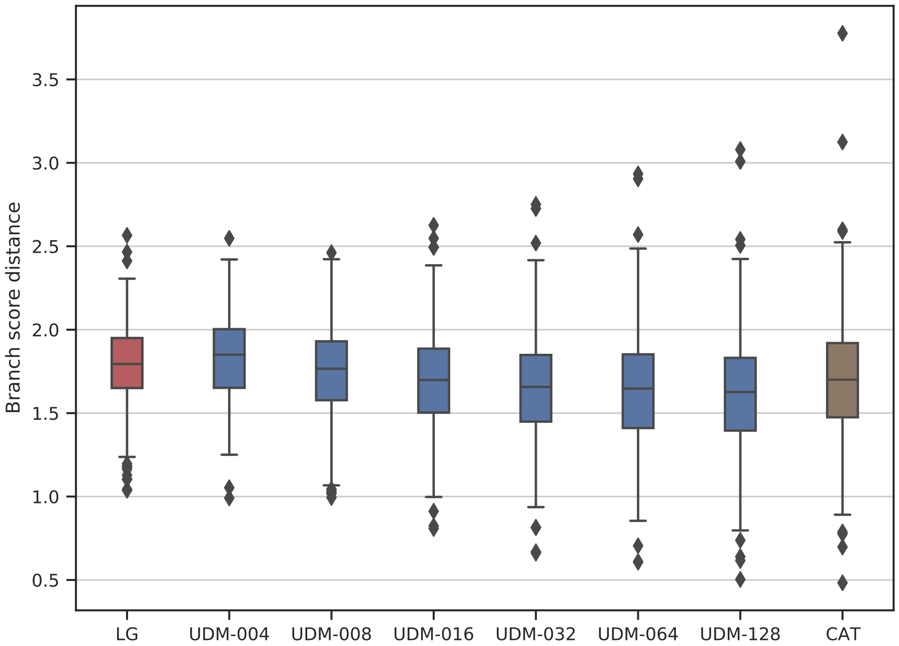

**Figure S17:** Branch score distances (Kuhner et al., 1994) obtained from the gene alignments of the microsporidia data set (Brinkmann et al., 2005) when compared to the currently accepted tree T2 (Figure 3). Results for the LG model (Le et al., 2008), universal distribution mixture models with four to 128 components obtained from the log center log ratio (Godichon-Baggioni et al., 2018) transformed site distributions, and the CAT model (Lartillot et al., 2004) with Poisson exchangeabilities are shown.

### S6 Results for nematodes

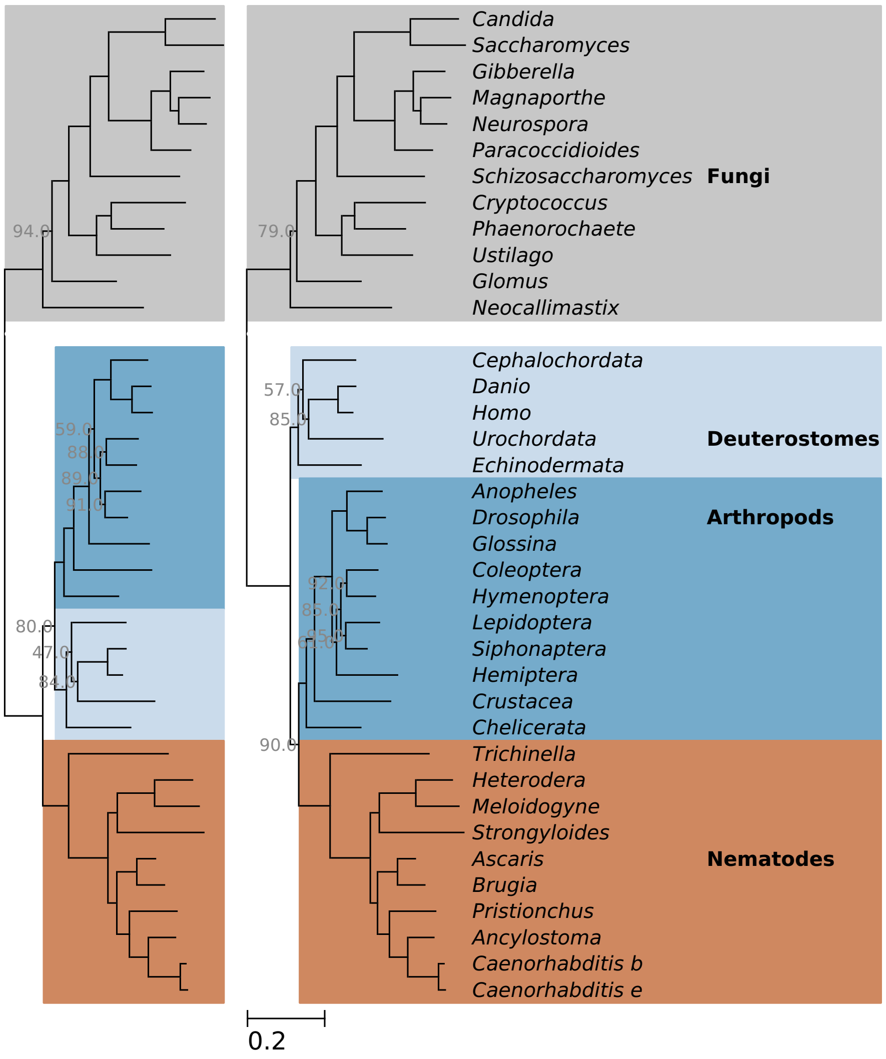

**Figure S18:** See Figure S10, but for nematode data set (Philippe et al., 2005). A universal distribution mixture model with only eight components was used on the right and side.

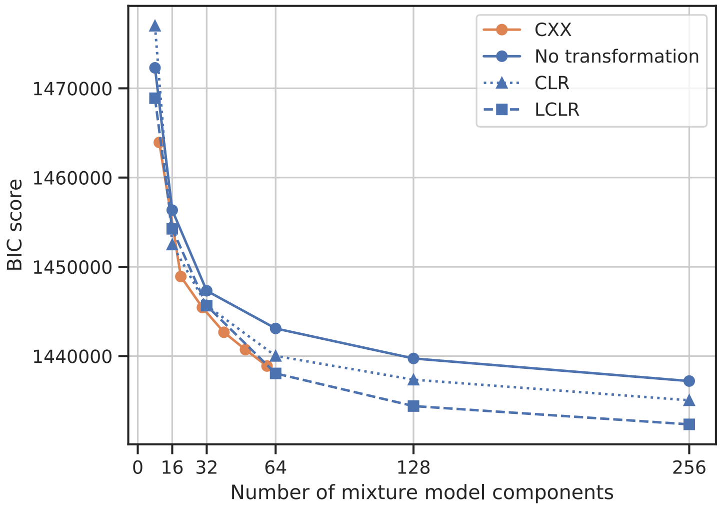

**Figure S19:** Bayesian information criterion (BIC; Schwarz, 1978) scores for different empirical distribution mixture models of the nematode data set (Philippe et al., 2005).

### S7 Results for platyhelminths

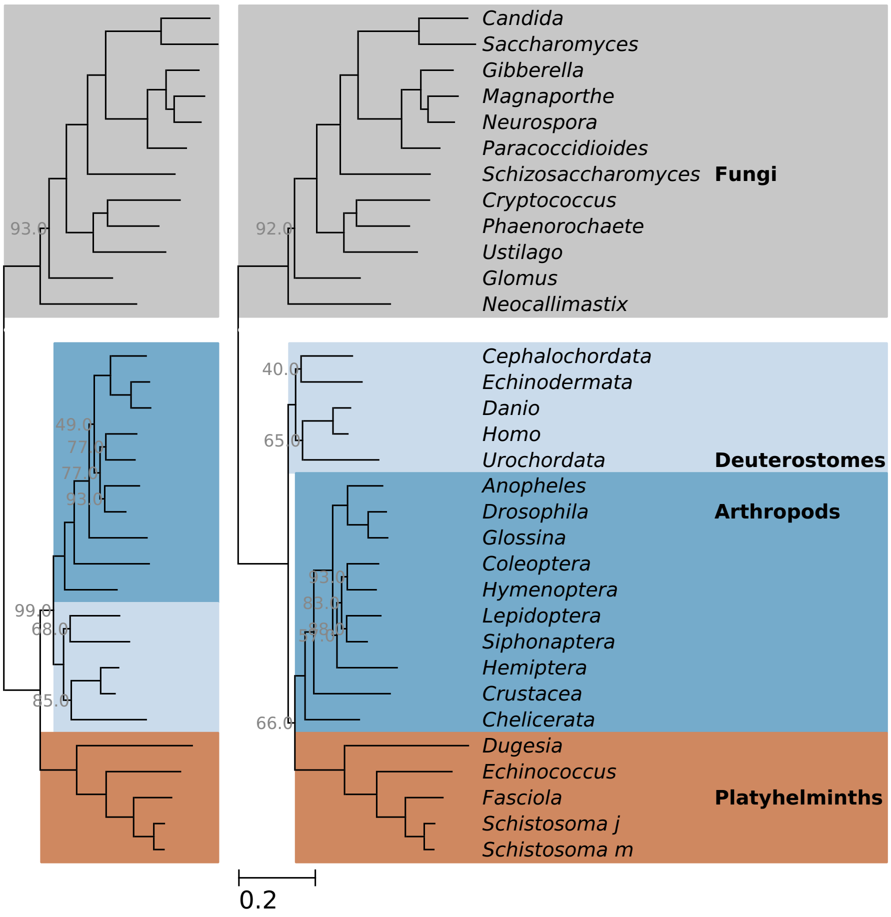

**Figure S20:** See Figure S10, but for platyhelminth data set (Philippe et al., 2005). A universal distribution mixture model with only 16 components was used on the right hand side.

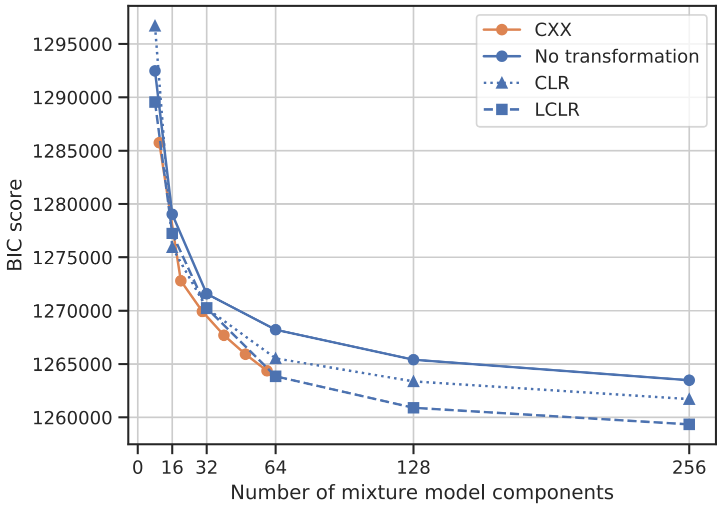

**Figure S21:** Bayesian information criterion (BIC; Schwarz, 1978) scores for different empirical distribution mixture models of the platyhelminth data set (Philippe et al., 2005).

### S8 Tool for phylogenetic analysis

During the course of this project, small tools for phylogenetic analysis were developed. The set of tools was collected in a software package termed Elynx Suite (`https://github.com/dschrempf/elynx`). The Elynx Suite is a Haskell library and a command line tool set for computational biology. The goal of the Elynx Suite is reproducible research. Evolutionary sequences and phylogenies can be read, viewed, modified and simulated. Thereby, the tools do no assume anything about the data, for example, about the type of code. No options are provided with default values. Consequently, the exact command with all arguments has to be stated by the user and is logged automatically, so that the analysis can be repeated easily. Usually, the initial work overhead is well compensated by the gain of reproducibility.

For example, to simulate a sequence of length 25 000 along a phylogeny TREE with a set of four stationary distributions DISTS (Phylobayes format; see Section S4), with custom weights and Poisson exchangeabilities, the following command line can be used:

```
slynx simulate -t TREE -e DISTS -w "[0.4415,0.2585,0.2178,0.0822]" \
-g "(4,0.8357)" -m "EDM(Poisson-Custom)" -l 25000
```

#### S9 Parametric bootstrap

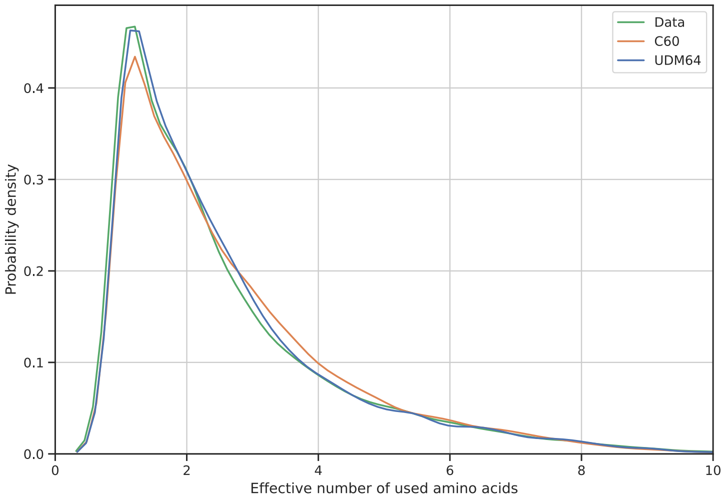

**Figure S22:** Kernel density estimation of the distribution of effective number of amino acids for the microsporidia data set (Brinkmann et al., 2005), and simulated data using the maximum likelihood estimates of the C60 (Quang et al., 2008), and the universal distribution mixture model with 64 components (UDM64) obtained from the log centered log ratio (Godichon-Baggioni et al., 2018) transformed site distributions of the HOGENOM database (Dufayard et al., 2005).

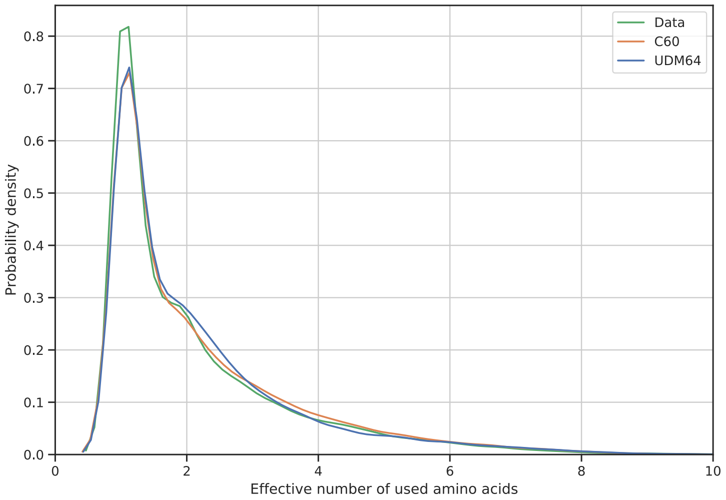

**Figure S23:** See Figure S22 but for the nematode data set (Philippe et al., 2005).

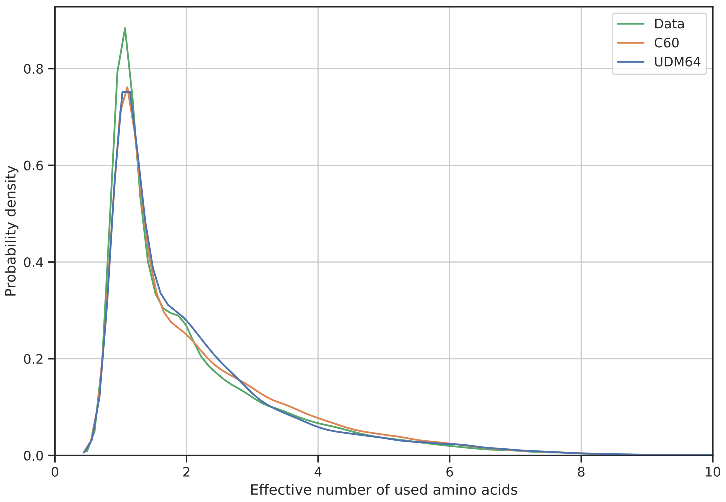

**Figure S24:** See Figure S22 but for the platyhelminth data set (Philippe et al., 2005).

| Model | $K$ | $r_{ij}$ | Transf. | $K_{\text{eff}}$ | log lh | $d_W$ |
|---|---|---|---|---|---|---|
| UDM | 256.0 | P | lclr | 2.555 | -719203.1 | 0.05 |
| Data | — | — | None | 2.568 | — | — |
| UDM | 256.0 | P | None | 2.577 | -720505.7 | 0.04 |
| UDM | 256.0 | P | clr | 2.588 | -719366.6 | 0.05 |
| UDM | 128.0 | P | lclr | 2.590 | -721134.7 | 0.03 |
| UDM | 64.0 | P | lclr | 2.592 | -723367.2 | 0.04 |
| UDM | 128.0 | P | None | 2.598 | -722555.8 | 0.05 |
| UDM | 32.0 | P | lclr | 2.619 | -726786.2 | 0.06 |
| UDM | 32.0 | P | clr | 2.621 | -725431.8 | 0.08 |
| UDM | 128.0 | P | clr | 2.631 | -721125.8 | 0.08 |
| UDM | 64.0 | P | None | 2.637 | -724584.3 | 0.07 |
| UDM | 32.0 | P | None | 2.645 | -726836.7 | 0.09 |
| UDM | 64.0 | P | clr | 2.651 | -722771.3 | 0.08 |
| UDM | 16.0 | P | None | 2.651 | -731115.1 | 0.12 |
| CXX | 40.0 | P | None | 2.652 | -724397.8 | 0.10 |
| UDM | 16.0 | P | lclr | 2.663 | -731000.4 | 0.10 |
| CXX | 60.0 | P | None | 2.671 | -722924.3 | 0.11 |
| UDM | 16.0 | P | clr | 2.693 | -728966.2 | 0.12 |
| CXX | 20.0 | P | None | 2.693 | -727743.5 | 0.13 |
| CXX | 30.0 | P | None | 2.701 | -725786.2 | 0.14 |
| UDM | 8.0 | P | lclr | 2.727 | -738395.7 | 0.15 |
| CXX | 50.0 | P | None | 2.737 | -723603.1 | 0.17 |
| UDM | 8.0 | P | clr | 2.761 | -740494.1 | 0.20 |
| UDM | 8.0 | P | None | 2.770 | -738437.8 | 0.20 |
| CXX | 10.0 | P | None | 2.779 | -735070.9 | 0.21 |
| UDM | 4.0 | P | clr | 2.845 | -754153.5 | 0.27 |
| UDM | 4.0 | P | lclr | 2.897 | -760962.9 | 0.32 |
| UDM | 4.0 | P | None | 2.911 | -763808.1 | 0.34 |
| WAG | 1.0 | WAG | None | 2.991 | -739218.5 | 0.42 |
| LG | 1.0 | LG | None | 3.037 | -731785.6 | 0.46 |

**Table S1:** Results of the parametric bootstrap analyses (see main text) of the microsporidia data set (Brinkmann et al., 2005). For each model, the average effective number of amino acids ($K_{\text{eff}}$) of the simulated data, the maximum log likelihood (log lh) obtained from the original data, and the Wasserstein distance ($d_W$) between the distributions of effective number of amino acids are given.

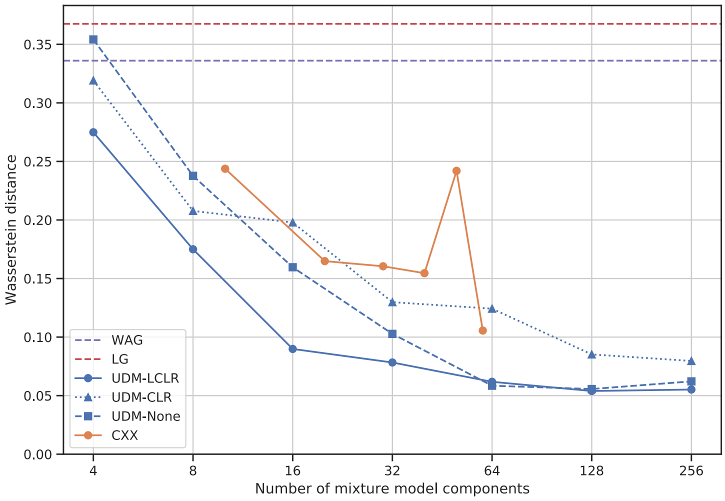

**Figure S25:** See Figure 4 in the main text but for the nematode data set (Philippe et al., 2005).

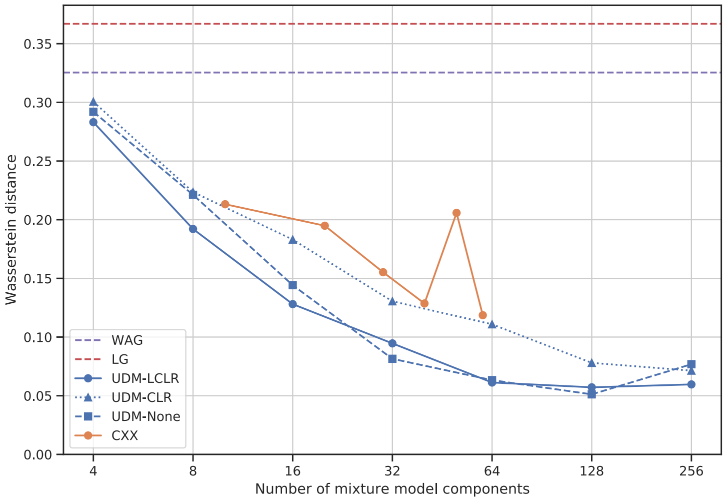

**Figure S26:** See Figure 4 in main text but for the platyhelminth data set (Philippe et al., 2005).

### S10 Simulation study

This section presents some additional results on the simulation study motivated by the microsporidia data set (Brinkmann et al., 2005). First, the LG model shows similar behavior than the Poisson model in that it infers the wrong topology (Figure S27). Please note the wrong position of the ciliates and that all branches are fully supported.

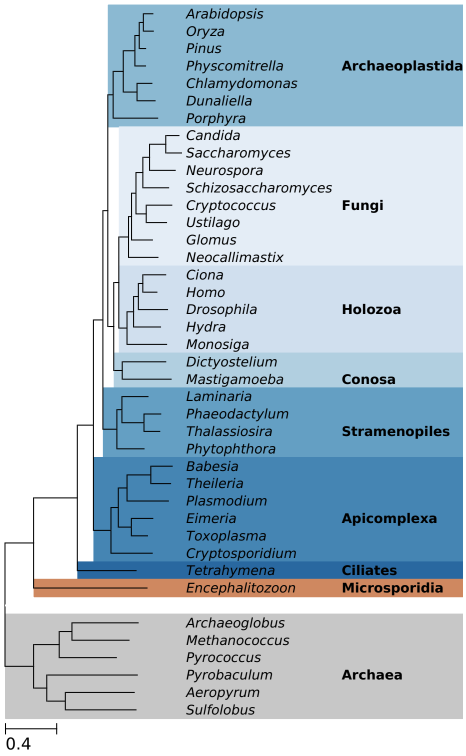

**Figure S27:** See Figure 5 in main text, but for the LG (Le et al., 2008) model.

Second, the C10 to C50 models (Quang et al., 2008), and the UDM models with eight and 16 components fail to infer the correct topology (Figure S28). The UDM model with 32 components inferred from the log center log ratio (LCLR, Godichon-Baggioni et al., 2018) transformed site distributions already infers the correct topology.

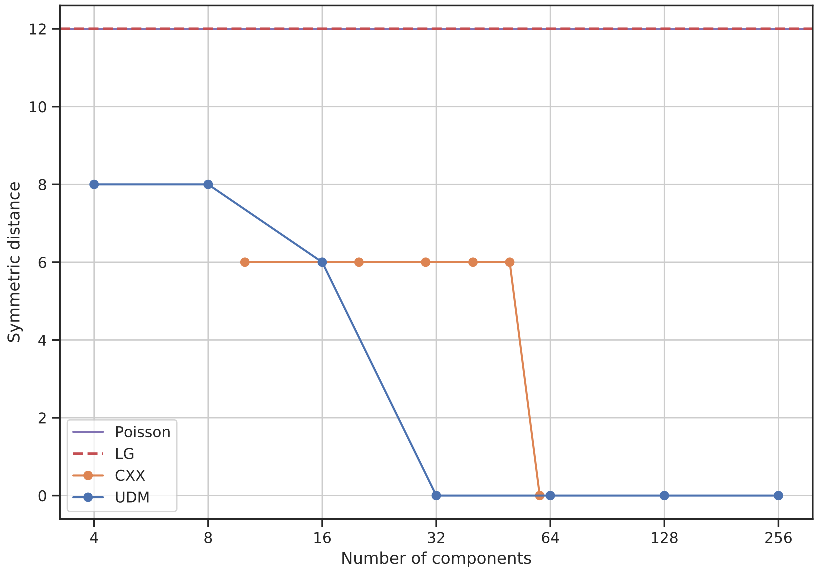

**Figure S28:** Symmetric distances (Robinson et al., 1981) for CXX (Quang et al., 2008) and universal distribution mixture (UDM) models with different numbers of components. The distances were calculated between the inferred phylogenies and original phylogeny used to simulate the alignment. For the UDM models, the log center log ratio transformation (Godichon-Baggioni et al., 2018) was used.

Further, the branch score distances (Kuhner et al., 1994) between the inferred phylogenies and the original phylogeny used to simulate the alignment show, at least partially, linear behavior in the $\log x$ plot (Figure S29). Consequently, the true value is approached by a function proportional to $\log N$, were $N$ is the number of components. The C60 model shows exceptionally good performance when compared to the C10 to C50 models because it is the first to infer the correct topology (Figure S28). However, the CXX models are still outperformed by the universal distribution mixture (UDM) models obtained from LCLR transformed site distributions.

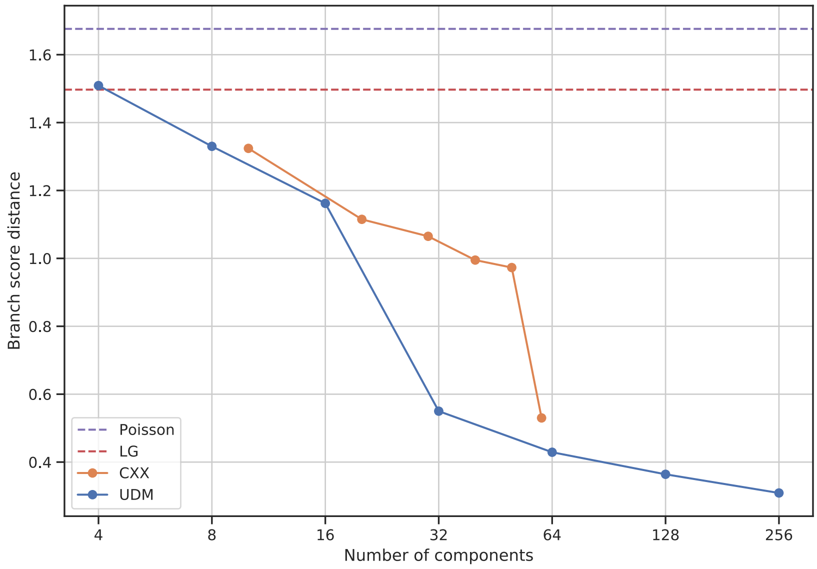

**Figure S29:** See Figure S28, but using the branch score distance.

Finally, the long alignment spanning 25 000 columns was randomly subsampled with replacement to provide a number of shorter alignments with lengths 100 to 1000 (Section 7). Also in these inferences, we observe that UDM models with many components outperform UDM models with fewer components, as well as classical substitution models such as the Poisson (Felsenstein, 1981) or the LG Le et al., 2008 models (Figures S30, and S31).

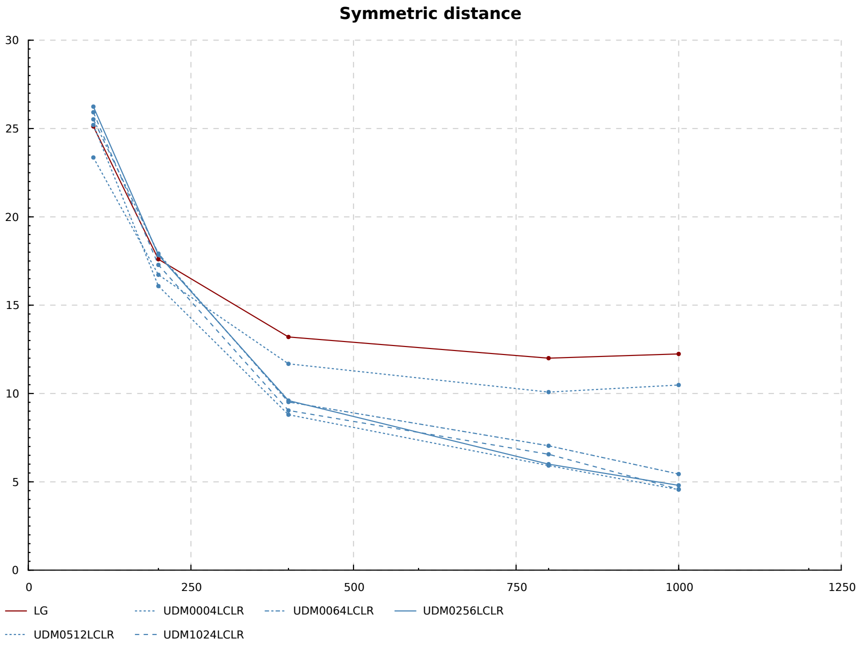

**Figure S30:** Averages of symmetric distances between trees inferred by the LG model (Le et al., 2008), as well as by various UDM models obtained from data transformed with the LCLR transformation (Godichon-Baggioni et al., 2018) and the original tree used for simulation. The x-axis traverses different alignment lengths.

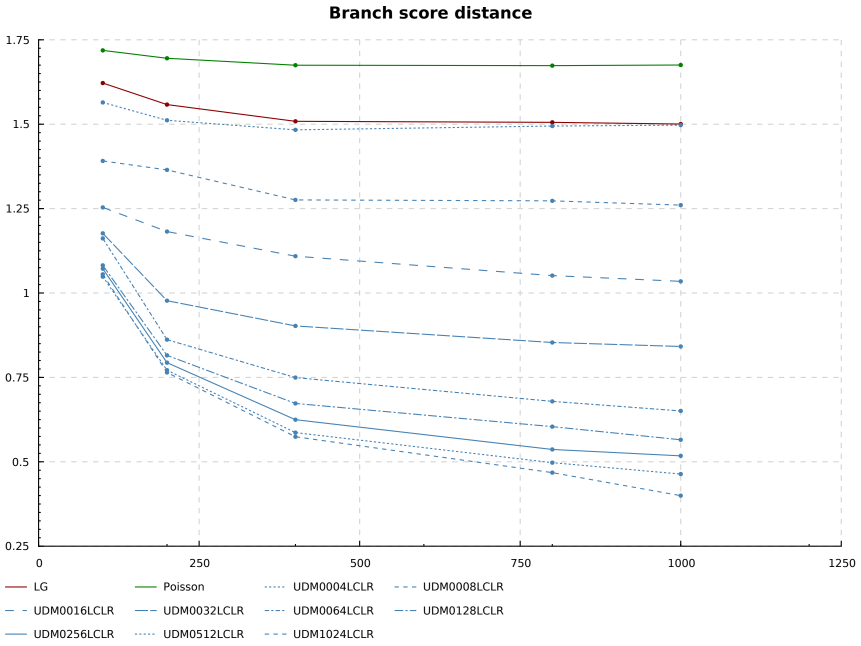

**Figure S31:** See Figure S30, but using the branch score distance.

#### S11 Divisive clustering

In order to reduce the redundancy between components of componen-rich UDM models, we tried a form of divisive clustering. Therefore, we start with one cluster constisting of all site distributions. Then, the cluster with the cluster center exhibiting the highest effective number of amino acids is divided into $K$ clusters (divisor) using $K$-means clustering. This is done repeatedly until the desired number of total clusters is reached. The results of phylogenetic inferences using UDM models obtained by divisive clustering are promising but not significantly different to the results of UDM models obtained by simple $K$-means clustering (Figures S32, and S33).

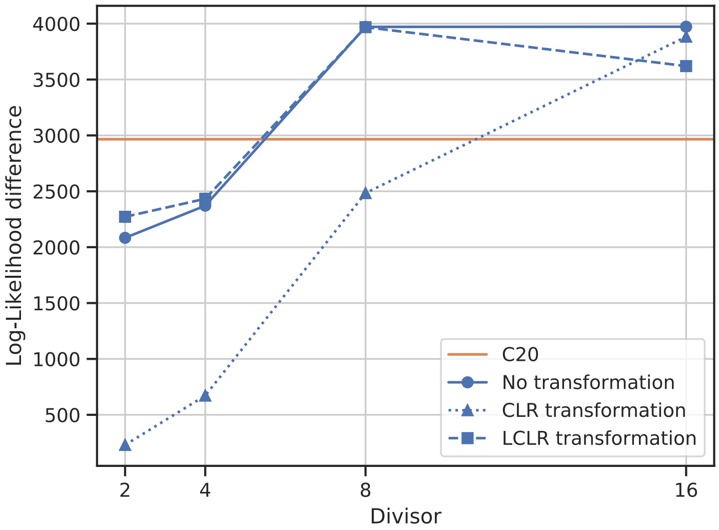

**Figure S32:** Log-likelihood difference between phylogenetic inferences using universal distribution mixture (UDM) models constrained to topologies T1 and T2 of the microsporidia data set (Brinkmann et al., 2005), respectively (see Figure 3 in main text). UDM models with 16 components obtained by divisive clustering using different divisors are used. For comparison, the result of the C20 model (Quang et al., 2008) is shown.

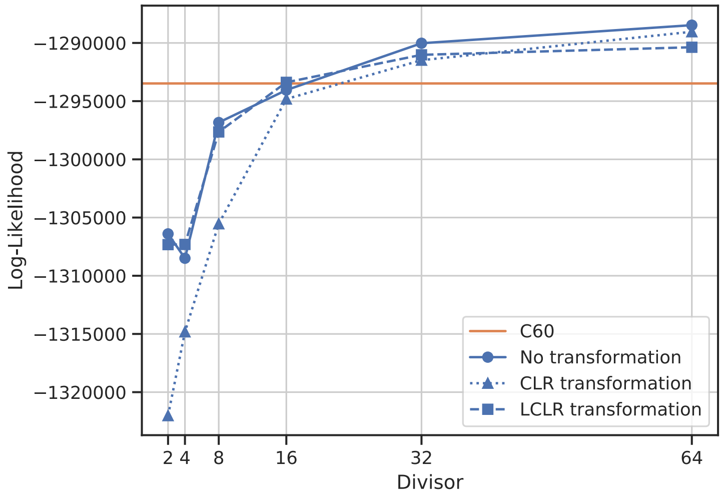

**Figure S33:** See Figure S32 but for 64 components. For comparison, the result of the C60 model (Quang et al., 2008) is shown.

### References

Aitchison J. 1982. The Statistical Analysis of Compositional Data. *J. Royal Stat. Soc. Ser. B (Methological).* 44:139–177.

Brinkmann H, Van Der Giezen M, Zhou Y, De Raucourt G P, Philippe H. 2005. An empirical assessment of long-branch attraction artefacts in deep eukaryotic phylogenomics. *Syst. Biol.* 54:743–757. DOI: 10.1080/10635150500234609.

Crooks G E. 2004. WebLogo: A Sequence Logo Generator. *Genome Res.* 14:1188–1190. DOI: 10.1101/gr.849004.

Dufayard J.-F, Duret L, Penel S, Gouy M, Rechenmann F, Perrière G. 2005. Tree pattern matching in phylogenetic trees: automatic search for orthologs or paralogs in homologous gene sequence databases. *Bioinformatics.* 21:2596–2603. DOI: 10.1093/bioinformatics/bti325.

Felsenstein J. 1981. Evolutionary trees from DNA sequences: a maximum likelihood approach. *J. Mol. Evol.* 17:368–376.

Godichon-Baggioni A, Maugis-Rabusseau C, Rau A. 2018. Clustering transformed compositional data using K-means, with applications in gene expression and bicycle sharing system data. *J. Appl. Stat.* 46:47–65. DOI: 10.1080/02664763.2018.1454894.

Höhna S, Landis M J, Heath T A, Boussau B, Lartillot N, Moore B R, Huelsenbeck J P, Ronquist F. 2016. RevBayes: Bayesian phylogenetic inference using graphical models and an interactive model-specification language. *Syst. Biol.* 65:726–736. DOI: 10.1093/sysbio/syw021.

Kuhner M K, Felsenstein J. 1994. A simulation comparison of phylogeny algorithms under equal and unequal evolutionary rates. *Mol. Biol. Evol.* 11:459–468.

Lartillot N, Philippe H. 2004. A Bayesian mixture model for across-site heterogeneities in the amino-acid replacement process. *Mol. Biol. Evol.* 21:1095–1109. DOI: 10.1093/molbev/msh112.

Lartillot N, Rodrigue N, Stubbs D, Richer J. 2013. PhyloBayes MPI: Phylogenetic Reconstruction with Infinite Mixtures of Profiles in a Parallel Environment. *Syst. Biol.* 62:611–615. DOI: 10.1093/sysbio/syt022.

Le S Q, Gascuel O. 2008. An improved general amino acid replacement matrix. *Mol. Biol. Evol.* 25:1307–1320. DOI: 10.1093/molbev/msn067.

Nguyen L.-T, Schmidt H A, Haeseler A von, Minh B Q. 2015. IQ-TREE: A Fast and Effective Stochastic Algorithm for Estimating Maximum-Likelihood Phylogenies. *Mol. Biol. Evol.* 32:268–274. DOI: 10.1093/molbev/msu300.

Pedregosa F et al. 2011. Scikit-learn: Machine Learning in Python. *J. Mach. Learn. Res.* 12:2825–2830.

Philippe H, Lartillot N, Brinkmann H. 2005. Multigene analyses of bilaterian animals corroborate the monophyly of Ecdysozoa, Lophotrochozoa, and protostomia. *Mol. Biol. Evol.* 22:1246–1253. DOI: 10.1093/molbev/msi111.

Quang L S, Gascuel O, Lartillot N. 2008. Empirical profile mixture models for phylogenetic reconstruction. *Bioinformatics.* 24:2317–2323. DOI: 10.1093/bioinformatics/btn445.

Robinson D F, Foulds L R. 1981. Comparison of phylogenetic trees. *Math. Biosci.* 53:131–147. DOI: 10.1016/0025-5564(81)90043-2.

Schwarz G E. 1978. Estimating the Dimension of a Model. *The Ann. Stat.* 6:461–464. DOI: 10.1214/aos/1176344136.

Tavaré S. 1986. Some Probabilistic and Statistical Problems in the Analysis of DNA Sequences. *Lect. on Math. Life Sci.* 17:57–86.

Whelan S, Goldman N. 2001a. A general empirical model of protein evolution derived from multiple protein families using a maximum-likelihood approach. *Mol. Biol. Evol.* 18:691–699. DOI: 10.1093/oxfordjournals.molbev.a003851.

Whelan S, Liò P, Goldman N. 2001b. Molecular phylogenetics: state-of-the-art methods for looking into the past. *Trends Genet.* 17:262–272. DOI: 10.1016/S0168-9525(01)02272-7.
